# A common neural currency account for social and non-social decisions

**DOI:** 10.1101/2021.10.18.464762

**Authors:** Desislava H. Arabadzhiyska, Oliver G. B. Garrod, Elsa Fouragnan, Emanuele De Luca, Philippe G. Schyns, Marios G. Philiastides

**Affiliations:** School of Psychology and Neuroscience, University of Glasgow, Glasgow, UK; Centre for Cognitive Neuroimaging, University of Glasgow, Glasgow, UK; School of Psychology, University of Plymouth, Plymouth, UK

## Abstract

To date, social and non-social decisions have been studied in isolation. Consequently, the extent to which social and non-social forms of decision uncertainty are integrated using shared neurocomputational resources remains elusive. Here, we address this question using simultaneous EEG-fMRI and a task in which decision evidence in social and non-social contexts varies along comparable scales. First, we identify comparable time-resolved build-up of activity in the EEG, akin to a process of evidence accumulation. We then use the endogenous trial-by-trial variability in the slopes of these accumulating signals to construct parametric fMRI predictors. We show that a region of the posterior-medial frontal cortex (pMFC) uniquely explains trial-wise variability in the process of evidence accumulation in both the social and non-social contexts. We further demonstrate a task-dependent coupling between the pMFC and regions of the human valuation system in dorso- and ventro-medial prefrontal cortex (dmPFC/vmPFC) across both contexts. Finally, we report domain-specific representations in regions known to encode the early decision evidence for each context. These results are suggestive of a domain-general decision-making architecture, whereupon domain-specific information is likely converted into a “common currency” in the dmPFC/vmPFC and accumulated for the decision in the pMFC.

## Introduction

Most strategic decisions occur under considerable uncertainty. For example, when investing in the stock market, a trader may use only purely probabilistic models to estimate risk in the market’s fluctuations. In contrast, when negotiating a deal in person, the trader’s risk assessment may rely instead on how trustworthy the other party appears (Fouragnan et al., 2013; Griessinger and Coricelli, 2015). Similarly, the decision to undergo a risky surgical operation may use online statistics regarding overall success rates or the advice of a trustworthy person who has undergone a similar operation.

In standard economic utility models (Morgenstern and Von Neumann, 1953), the rules governing such decisions are the same, regardless of whether the source of uncertainty is social or non-social in nature (e.g. a human or an on-line platform). Recent advances in social neuroscience, however, have focused instead on identifying neurocognitive processes that might be uniquely social (Rilling, King-Casas, and Sanfey, 2008; Suzuki and O’Doherty, 2020; van Baar, Chang, and Sanfey, 2019). Correspondingly, two competing accounts have recently been proposed to explain how the brain might encode uncertainty representations underlying social and non-social choices (Ruff and Fehr, 2014).

The first account assumes that the brain dedicates largely separate networks for encoding social and non-social forms of decision uncertainty and for assigning value to different choice alternatives. In contrast, the second account proposes that the same network processes the different forms of uncertainty and converts the values associated with different choice alternatives into a “common currency”. In order to arbitrate between these accounts, both the algorithmic (i.e. what computational mechanisms are involved) and implementational (i.e. which brain regions are involved) levels need to be considered (Lockwood, Apps, and Chang, 2020).

To date, however, there is no unified framework for integrating social and non-social decisions as most studies have evaluated them in isolation. Studies of non-social decision-making have focused primarily on constructing a mechanistic account of choice behavior using sequential sampling models in which evidence accumulates stochastically to an internal decision boundary (Hu, Domenech, and Pessiglione, 2020; Hunt et al., 2012; Sepulveda et al., 2020). Recent brain imaging work has implicated the posterior-medial frontal cortex (pMFC) in this process of evidence accumulation, proposing a domain-general role for this region in perceptual and value-based choices (Cona, Marino, and Semenza, 2017; Pisauro, Fouragnan, Retzler, and Philiastides, 2017; Zhang, Hughes, and Rowe, 2012).

While some recent work on social decision making started to explore the utility of such accumulation-to-bound models (Chen and Krajbich, 2018; Krajbich et al., 2015; Suzuki and O’Doherty, 2020) no *direct* comparisons have been made between the mechanisms and neural representations of social and non-social choices. Here, we design a novel task in which decision evidence in social and non-social contexts varies along comparable scales. Across contexts, we test whether there is a common embedding of decision evidence as well as a common mechanism for integrating this evidence using simultaneous electroencephalography (EEG) and functional magnetic resonance imaging (fMRI) (henceforth EEG-fMRI).

In doing so, we identify in both contexts centroparietal EEG signals exhibiting decision dynamics consistent with a common process of evidence accumulation. Consistent with such domain-general mechanisms, the trial-by-trial temporal variability in these accumulating signals is reflected in the fMRI data in the region of the pMFC previously implicated in other types of decisions (Pisauro, Fouragnan, Retzler, and Philiastides, 2017). Moreover we report a trial-wise and task-dependent modulation of the pMFC with established regions of the human valuation system in the medial prefrontal cortex (Chib, Rangel, Shimojo, and O’Doherty, 2009; Clithero and Rangel, 2014; Philiastides, Biele, and Heekeren, 2010), suggestive of decision evidence embedded within a “common currency” space.

## Results

We investigated economic decisions within a social context by exploiting trustworthiness in a partner’s face during a strategic economic game to generate predictions about possible outcomes (see below) and within a non-social (purely probabilistic) context by manipulating outcome probabilities in individual gambles. Importantly, we created parametrically modulated stimuli along comparable scales of reward probability in each of the social and non-social contexts (while keeping reward magnitude constant across both contexts). The non-social stimuli were associated with a range of pure reward probabilities chosen from the full probability range (from 0–1), placed on top of a face image (neutral for trustworthiness) to equalize perceptual load across the social and non-social stimuli (Fig. 1; see Materials and Methods for more details).

**Figure 1.**
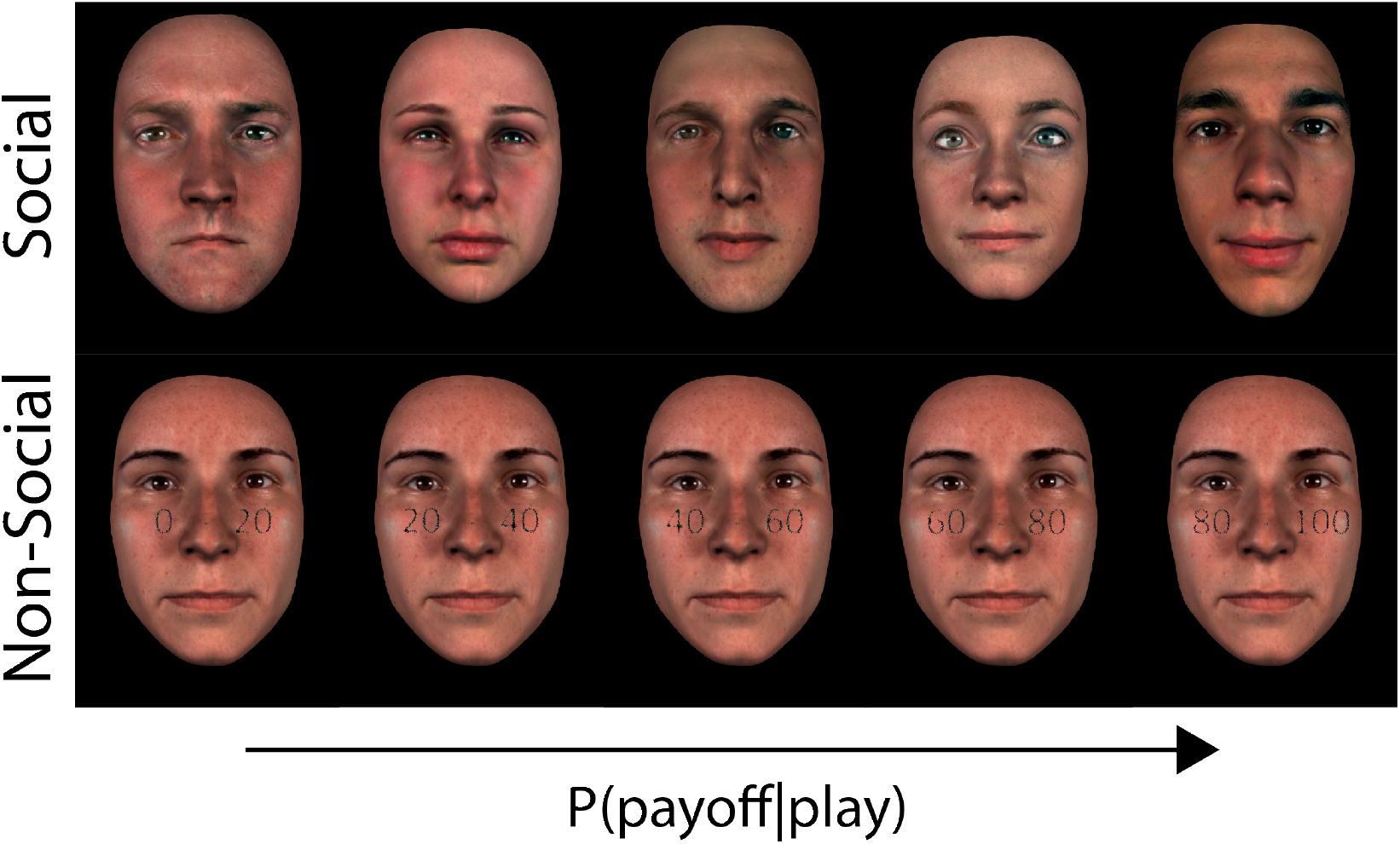
Sample stimuli from a representative participant. Top: Social stimuli at five different (participant–specific) indirect trustworthiness levels, matching the pure reward probability levels used for the non-social stimuli. For each participant there were, on average, 28 unique face identities in each of the five reward probability levels. Bottom: Non-social stimuli with five explicit reward probability levels (given a ’Play’ choice) superimposed on a face neutral for trustworthiness (i.e. 0.5 reward probability). Photo-realistic face images were obtained using the procedure described in (Gill, Garrod, Jack, and Schyns, 2014) and summarised in Materials and Methods.

We derived comparable reward probabilities for the social stimuli by asking participants (N=31) to provide indirect trustworthiness ratings for a series of 150 face identities. Specifically, we framed this rating stage in the context of a trust game (Berg, n.d.). Usually a trust game involves an interaction between two players, the investor and the trustee. The investor decides whether to send a monetary endowment to the trustee that gets multiplied by a certain factor (’Play’ option) or to retain possession of the initial endowment (’Keep’ option). In turn, the trustee can decide whether or not to send a fixed share of the augmented amount back to the investor so that both parties can benefit from the interaction. We told participants that each face belonged to people who had previously taken part in a similar study in the role of the trustee and we asked them to indicate the overall likelihood (in the range 0–1) that each person had returned a fixed share (50%) of the augmented endowment entrusted to them (Fig. 2a).

**Figure 2.**
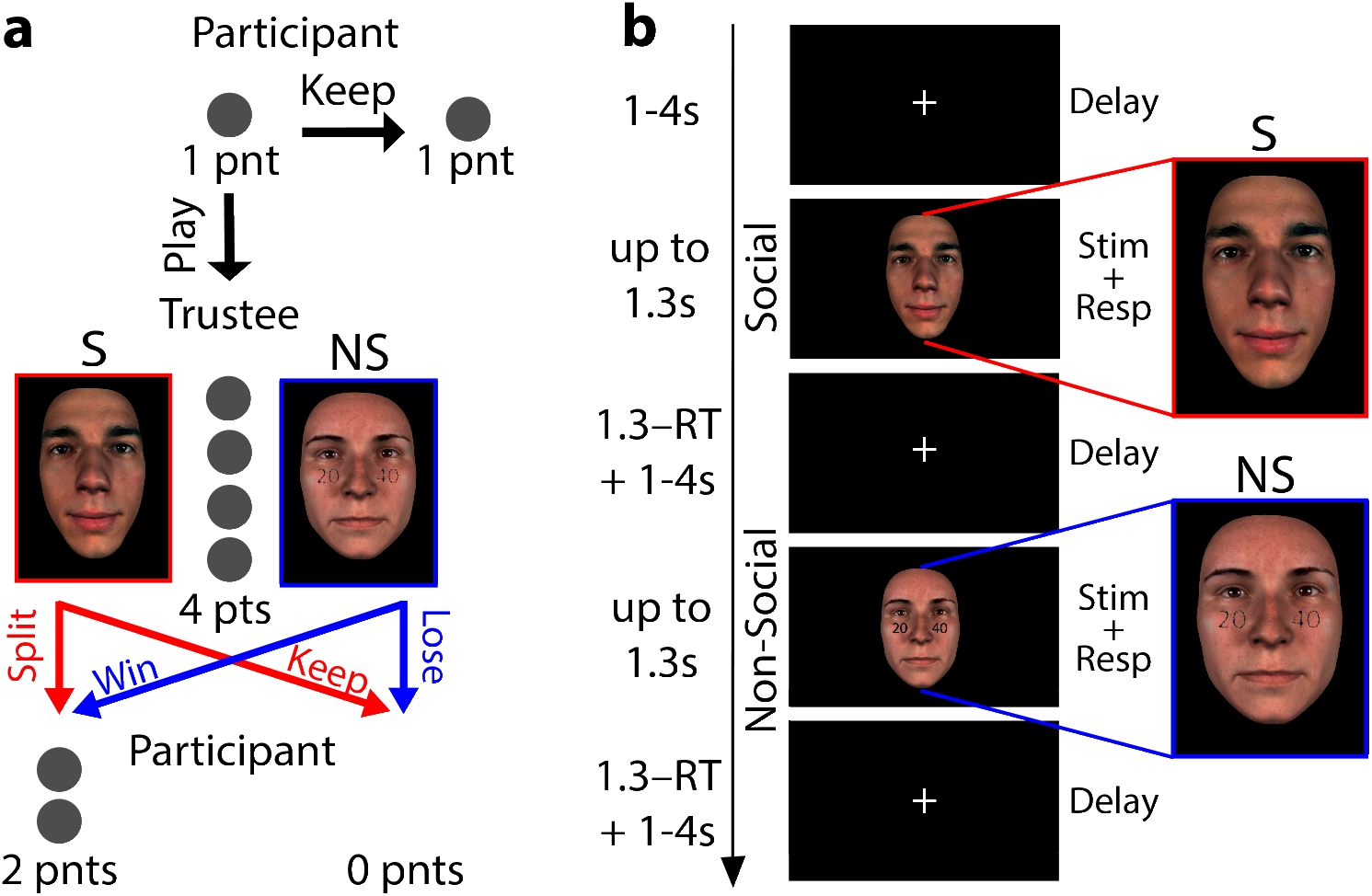
**a**. A variant of the traditional Trust Game in which a participant (Investor) is allocated 1 point and they need to decide whether to ’Keep’ the point or ’Play’ for the chance of winning 2 points. During ’Play’ choices the 1 point is quadrupled and passed on to a Trustee, which takes the form of either a social agent (red) or a purely probabilistic gamble (blue). The Trustee can either split the 4 points evenly and give the participant 2 points or keep all 4 points to themselves (i.e. the participants receives 0 points). In the social context the probability of winning is based on the trustworthiness of the social agent displayed in the stimulus, while in the non-social context by the reward probability range displayed on top of a face, neutral for trustworthiness. **b**. Social (S; red outline) and non-social (NS; blue-outline) experimental design trials. Each trial began with a variable duration (1-4 s) fixation cross screen, which served as an inter-trial interval. The fixation screen was followed by a stimulus screen which remained available for up to 1.3 s, during which participants indicated their choice (’Play’ or ’Keep’). The stimulus screen was replaced by a fixation cross following choice for the remainder of the 1.3 s.

This indirect measure of perceived trustworthiness was previously shown to be more ecologically valid compared to explicit ratings (Uleman and Kressel, 2013) and further ensured that trustworthiness judgments became the product of an economic decision as in our main experimental paradigm. Specifically, during the main (EEG-fMRI) task participants assumed the role of the investor in a series of one-shot trust games. In each game they had to decide whether to choose between a small but sure reward (1 pt; ’Keep’ option) or a bigger, but riskier payoff (2 pts; ’Play’ option). We randomly interleaved non-social trials (i.e. probabilistic gambles) in which we controlled the likelihood of obtaining the higher payoff with explicit reward probabilities, matched against subject-specific indirect trustworthiness ratings as highlighted above (Fig. 2b).

We instructed participants that the probability of receiving the higher payoff for ’Play’ choices would be based on the overall likelihood with which each face identify split the augmented endowment (here 4 pts) in a previous study (social trials) or the pure reward probabilities depicted on the face stimuli (non-social trials). We sampled the full range of reward probabilities given a ’Play’ choice using five levels (*P*(*payoff* |*play*) = {0 – 0.2,0.2 – 0.4,0.4 – 0.6,0.6 – 0.8,0.8 – 1}). In social trials, we populated each reward probability level with face identities based on the subject-specific perceived trustworthiness from the initial rating stage. Ultimately, these ranges correspond to three broad task difficulty levels; easier trials favouring either a ’Keep’ or ’Play’ choice (i.e. 0–0.2 and 0.8–1, respectively), medium difficulty trials in which the outcome uncertainty for ’Play’ choices begins to increase (i.e. 0.2–0.4 and 0.6–0.8) and difficult trials for the most ambiguous set of reward probabilities (i.e. 0.4–0.6).

### Behavioral performance in social and non-social contexts

Participants’ fraction of ’Play’ choices correlated positively with the overall reward probability across both the social and non-social trials (Social: t(30) = 17.769, p <0.001; Non-social: t(30) = 4.086 p <0.001), indicating that they selected the riskier option more frequently as the likelihood of receiving the higher payoff increased (Fig. 3a). More importantly, we demonstrated that choice behavior was comparable across the social and non-social trials. Specifically, we used a likelihood-ratio test (see Material and Methods) to show that a single sigmoid function fit the fraction of ’Play’ choices (jointly across both conditions) as well as two separate functions (*λ*(30) = 0.551, p = 0.759).

**Figure 3.**
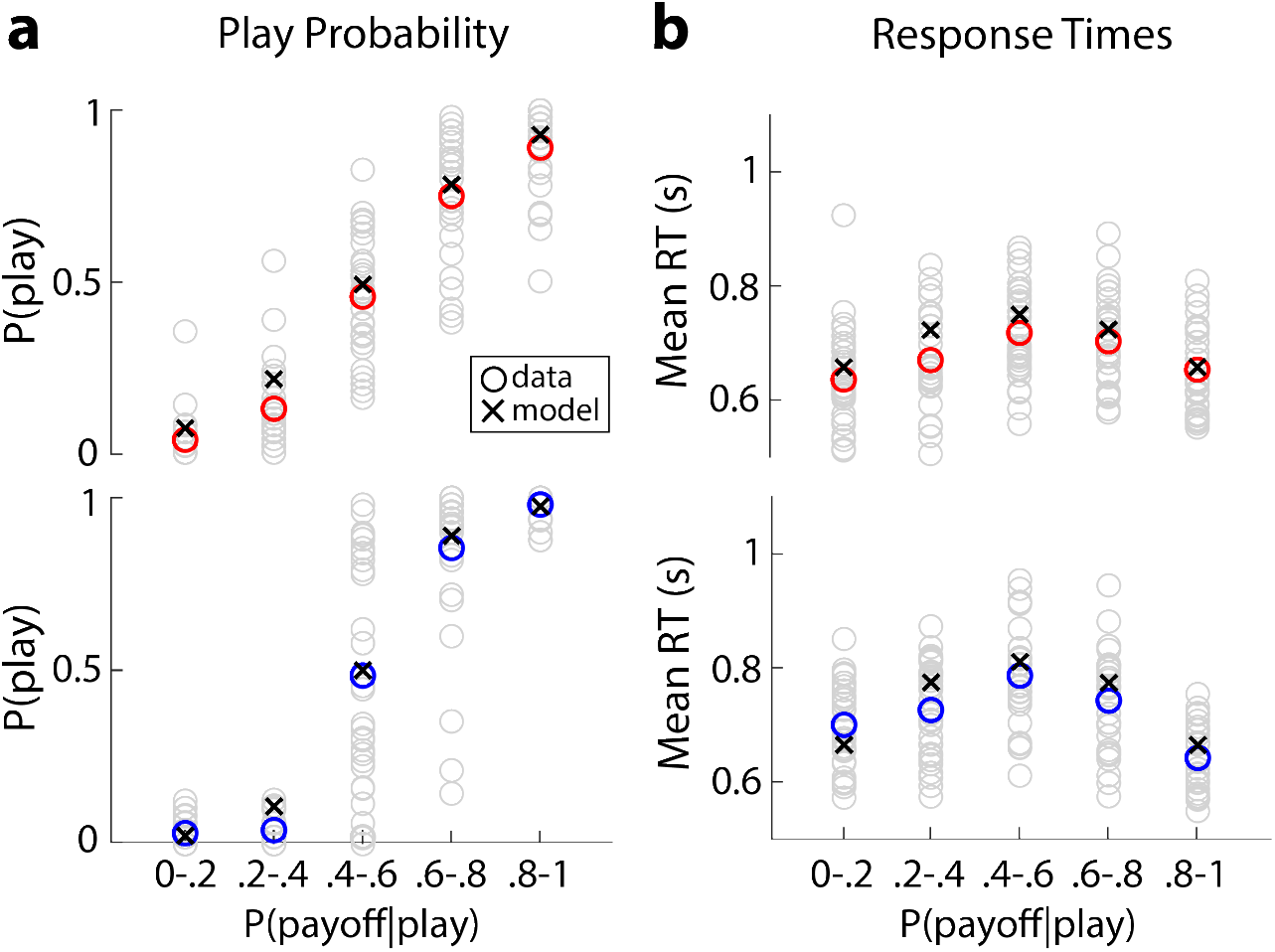
Social and non-social behavioral responses (red and blue circles) versus modelling performance of a drift diffusion model (black crosses) for proportion of ’Play’ choices (**b**) and response times (RTs; **c**). ’Play’ responses increased with probability of reward given a ’Play’ choice (P(payoff|play)) and RTs were the highest when there was no strong evidence for or against ‘Play’ decisions. Participant-specific behavior presented in grey circles.

The mean response times (RTs) as a function of the overall reward probability exhibited an inverted ‘V’ relationship, across both the social and non-social trials (Fig. 3b), consistent with a positive relationship with task difficulty (Social: t(30) = 9.302, p <0.001; Non-social: t(30) = 10.105, p <0.001). In other words, we observed the longest RTs for the most difficult trials (reward probabilities 0.4–0.6), the shortest RTs for the easiest trials (reward probabilities 0–0.2 and 0.8–1) and intermediate RTs for medium difficulty trials (reward probabilities 0.4–0.6 and 0.6–0.8). The overall RTs showed a small (41.637ms), albeit significant difference between the social and non-social trials (paired ttest: t(30) = −3.274, p = 0.003), with social trials (*M_S_* = 677.864ms, *SD_S_* = 86.479 ms) being on average faster than non-social ones (*M_NS_* = 719.502ms, *SD_NS_* = 91.287 ms).

### Evidence accumulation in social and non-social contexts

Having established comparable behavioral performance across the social and non-social contexts, we asked whether these share a common underlying mechanism for integrating relevant decision evidence. To address this question, we first leveraged the high temporal resolution of the EEG data to identify signals exhibiting a gradual build-up of activity consistent with a general process of evidence accumulation (EA) (Pisauro, Fouragnan, Retzler, and Philiastides, 2017; Polaniía, Krajbich, Grueschow, and Ruff, 2014). We hypothesize that if such signals exist, we should observe reliable ramp-like activity with a build-up rate that is proportional to the amount of decision difficulty.

We tested this hypothesis with a single-trial multivariate linear classifier (Parra, Spence, Gerson, and Sajda, 2005; Sajda, Philiastides, and Parra, 2009) designed to estimate spatial weightings of the EEG sensors that discriminate between easy vs. difficult trials (see Materials and Methods). Applying the estimated electrode weights to single-trial data produced a measurement of the discriminating component amplitudes (henceforth **y**). These amplitudes represent the distance of individual trials from the discriminating hyperplane that we treat as a neural surrogate for the relevant decision variable that is being integrated.

Persistent accumulating activity with a build-up rate proportional to the amount of decision difficulty should result in a gradual increase in the classifier’s performance while the traces for the easy and difficult trials diverge as a function of elapsed time in stimulus-locked data (Fig. 4a). To test the extent of a domain-general EA process across social and non-social contexts, we performed this discrimination by initially collapsing trials across both conditions. We treated the medium difficulty trials as “unseen” data (independent of those used to train the classifier), to more convincingly test for a full parametric effect on the build-up rate associated with the different levels of decision difficulty.

**Figure 4.**
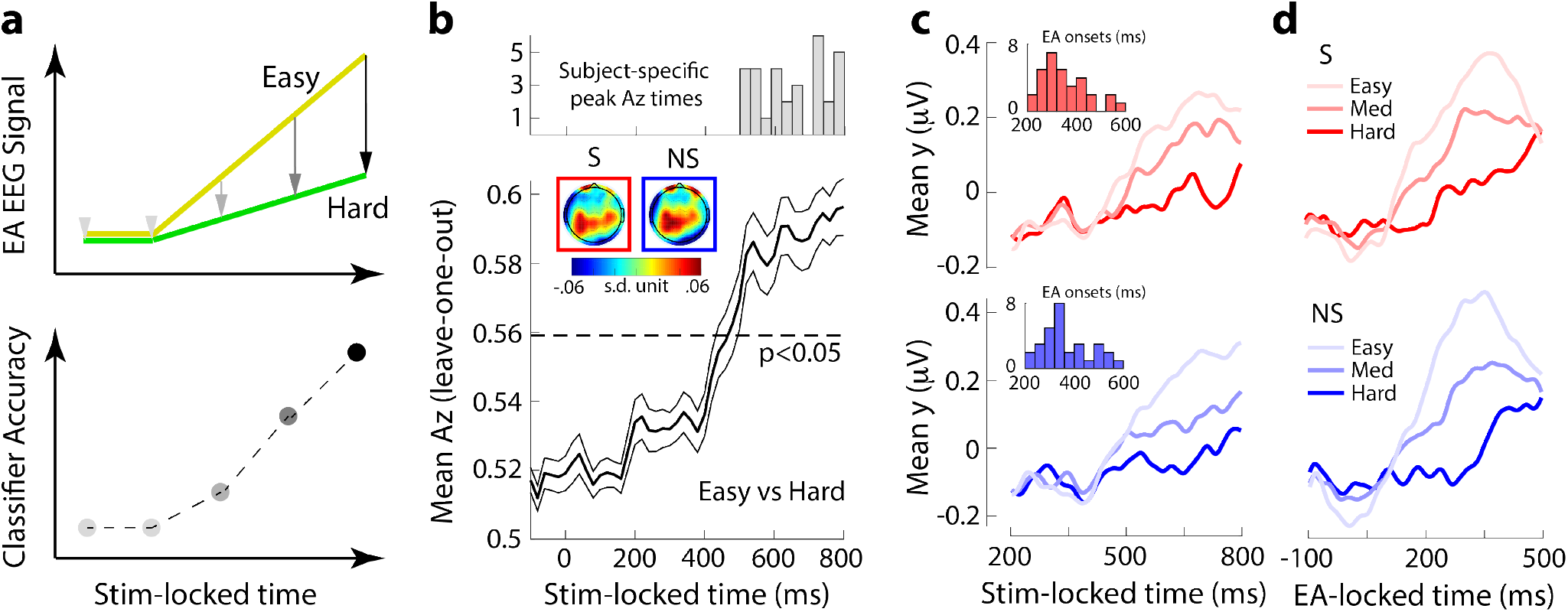
Linear discriminant analysis of the EEG. **a**. Build-up rates for hypothetical evidence accumulation (EA) signals for easy (yellow) and difficult (green) trials (top) and how differences in the rate of EA could manifest in the accuracy of an EEG classifier trained on stimulus-locked data. **b**. Average discrimination performance (Az; using leave-one-out cross validation) between easy and difficult trials across participants along with histogram of participant-specific peak discrimination times (top). The dashed line represents the average Az value leading to a significance level of p=0.05, estimated using a separate bootstrap test. The thinner black lines indicate standard errors of the mean across participants. Insets: scalp topographies (forward models) of the discriminating activity estimated at time of maximum discrimination averaged across participant for the social (red outline) and non-social (blue outline) trials. **c**. The average temporal profile of the discriminating activity across participants (obtained by applying the participant-specific classification weights estimated at the time of maximum discrimination) for the three levels of decision difficulty for social (red) and non-social trials (blue), locked to the onset of the stimulus onset. Insets: histograms of participant-specific EA onset times for social (red) and non-social trials (blue). **d**. The average temporal profile of the discriminating activity across participants, realigned to the onset of EA as estimated in **c**, for the three levels of decision difficulty for social (red) and non-social trials (blue).

We quantified the classifier’s performance through the area under a ROC curve (i.e. Az value) with a leave-one-trial-out cross validation procedure. This was done at several time windows locked to the stimulus onset. As hypothesized, the classifier’s performance increased systematically over time, reflecting the potential divergence in the gradual build-up of activity between easy and difficult trials (Fig. 4b). On average, the classifier’s performance began increasing after 400 ms post-stimulus (i.e. after early encoding of the relevant evidence) and peaked several hundred milliseconds later.

The spatial distribution of this discriminating activity (i.e. forward model; see Material and Methods) from participant-specific windows of maximum discrimination between easy and difficult trials (Fig. 4b, top) revealed comparable centroparietal topographies across social and non-social contexts (r = 0.896, p < 0.01; Fig. 4b, inset). These similarities are suggestive of common neural generators across the two contexts, consistent with those reported previously in the perceptual domain (e.g. Herding et al., 2019; Kelly and O’Connell, 2013; Philiastides, Heekeren, and Sajda, 2014).

To formally characterize the temporal profile of the discriminating activity (i.e. **y**(**t**)) for each condition separately, we applied participant-specific spatial weights from the time window of maximum discrimination to an extended stimulus-locked time window and separately for social and non-social trials. We also applied this procedure to medium difficulty trials (i.e., “unseen” data) by subjecting the relevant data through the same neural generators responsible for the original discrimination.

This approach revealed a gradual build-up of activity akin to a process of EA in both social and non-social trials (Fig. 4c; top: Social, bottom: Non-social). Similar to the classifier performance, the neural activity began to rise around 400 ms after stimulus presentation in both the social and non-social trials, with the build-up rate being proportional to the amount of decision difficulty. Note that the build-up rate from medium difficulty trials was situated between the two extreme conditions used to originally train the classifier, thereby establishing a fully parametric effect across the three levels of decision difficulty (F(2, 90)=16.88, p<0.001 for the social condition, F(2, 90) = 26.76, p<0.001 for the non-social condition, post-hoc paired t-tests, all p< 0.001).

We finally extracted participant-specific EA onset times, that is the time point at which the discriminating activity began to rise monotonically after an initial dip in the data following any early evoked responses present in the data (Fig. 4c; insets). Due to the inter-individual variability in these onset times, we predicted that re-aligning the relevant signals to the participant-specific EA onset times should reveal a more pronounced depiction of the underlying process of EA at the population level, which was indeed the case (Fig. 4d; top: Social, bottom: Non-social).

### Linking neural signatures of evidence accumulation to behavior

To further establish that our EEG signals reflect the process of EA leading up to the decision we performed two additional analyses to link these signals with our participants’ behavioral performance. Firstly, we expected the build-up rate of the EEG EA signals to correlate with drift rate estimates obtained from a drift diffusion model (DDM) (Pisauro, Fouragnan, Retzler, and Philiastides, 2017; Polaniía, Krajbich, Grueschow, and Ruff, 2014) fit on participants’ fraction of ’Play’ choices and RTs (Fig. 3; Social – Fraction ’Play’ Choice: r = 0.945, t(154) = 36.464, p<0.001, RT: r = 0.754; t(154) = 15.154, p<0.001; Non-social – Fraction ’Play’ Choice: r = 0.968, t(154) = 94.196, p<0.001, RT: r = 0.765; t(154) = 14.461 p<0.001).

We set the decision thresholds for ’Play’ and ’Keep’ choices to +1 and −1 respectively (see Materials and Methods for details). Correspondingly, positive (negative) drift rates are associated with reward probability levels favoring a ’Play’ (’Keep’) choice. To align the EEG build-up rates (i.e. linear slopes of **y**(**t**); see Materials and Methods) with this convention, we flipped the sign of the EEG slopes in the two reward probability levels which support ’Keep’ choices (i.e. *P*(*payoff* |*play*) = {0 – 0.2,0.2 – 0.4}). As expected, we observed robust correlations between the slopes of the EEG and drift rates, across both social (Fig. 5a; r = 0.653, p < 0.001) and non-social trials (Fig. 5b; r = 0.709, p < 0.001).

**Figure 5.**
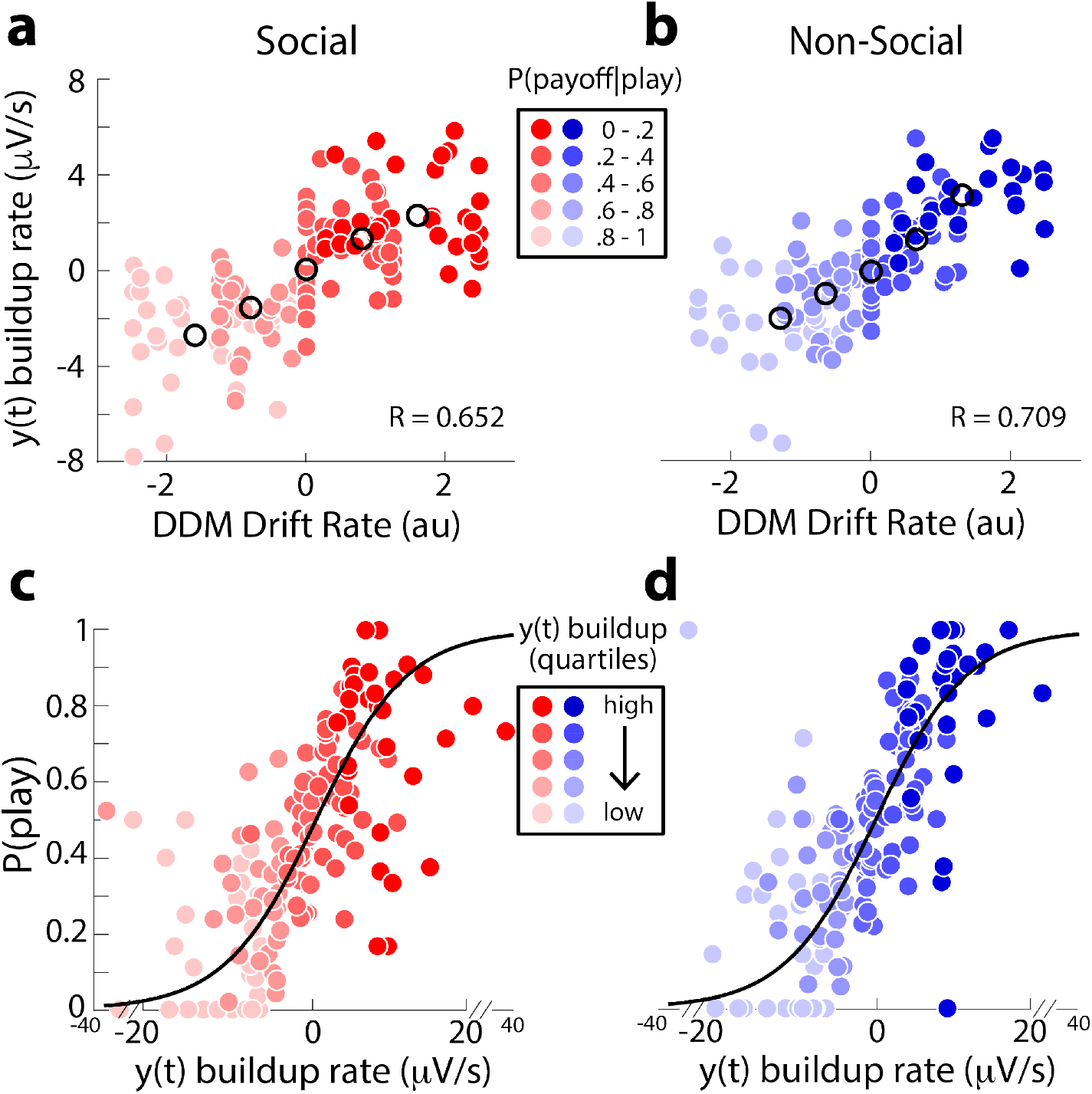
Linking EEG signatures of evidence accumulation to behavior. **a, b**. Participant-specific EA slopes (**y**(**t**)) build-up rate) for each of the five levels of *P*(*payoff* | *play*) scale positively with DDM estimates of drift rate for both the social (**a**) and non-social (**b**) contexts. The EEG-derived EA signal **y**(**t**) from which the EA slopes were derived was normalized per trial to factor out any effects unrelated to the EA processing, such as attentional drifts. Black circles indicate population averages. **c, d**. Trial-by-trial estimates of EA slopes correlate positively with the probability of playing (Eq.3) for both the social (**c**) and non-social (**d** contexts. To visualize this association the data points were computed by grouping trials into five bins based on the EA slope estimates. Importantly, the black curves are derived from fits of Eq. 3 to individual trials.

Secondly, we used the trial-by-trial slope estimates from the EEG EA signals to directly predict the probability of playing on individual trials, using a logistic regression. Consistent with the previous analysis we flipped the sign of the EEG slopes in the two reward probability levels which support ’Keep’ choices. We expected, high positive and high negative EA rates to reflect easy ’Play’ and ’Keep’ choices respectively, with intermediate magnitude slopes reflecting medium difficulty choices and slopes near zero representing difficult choices. As expected, we found that EEG slopes were a significant predictor of the eventual probability of playing for both the social (Fig. 5c; t(30) = 7.582, p<0.001) and non-social trials (Fig. 5d; t(30) = 8.173, p<0.001).

Finally, we also tested the possibility that trial-by-trial modulations of the EEG slopes may have arisen merely due to fluctuations in attention (as it waxes and wanes in the course of the experiment). Specifically, we ran a linear serial autoregression model predicting the EEG-derived EA slope in the current trial from the slopes from the previous four trials, individually for all participants. We found that on average this only accounted for a very small portion of the overall variance in the EEG slopes (Social: R^2^ = 0.02, Non-social: R^2^ = 0.019), demonstrating the absence of a serial autocorrelation in slopes across neighbouring trials.

### EEG-informed fMRI of evidence accumulation

Our EEG analysis demonstrated that both the social and non-social choices display comparable EA dynamics, potentially suggestive of common underlying neural generators. Here, we aimed to exploit endogenous trial-by-trial variability in the slope of EA to create EEG-informed fMRI predictors to identify candidate regions for the process of EA in social and non-social contexts (Fig. 6a). We note that the EA slopes derived from the EEG were not highly correlated with individual RTs (Social: r = −0.297, Non-Social: r = −0.333). This is due to the high degree of inter-trial variability in the decision and motor planning stages, as has been demonstrated consistently in previous modelling and experimental studies (Philiastides, Heekeren, and Sajda, 2014; Ratcliff, Philiastides, and Sajda, 2009; Verdonck, Loossens, and Philiastides, 2021). As such, RTs do not constitute a major confounding factor but we nonetheless included separate nuisance RT predictors in our fMRI analysis (for each of the social and non-social contexts). Similarly, for each context, we included parametric predictors based on the experimentally-defined task difficulty (i.e. easy, medium, difficult trials) to ensure that any overall task-difficulty effects (e.g. in the slopes of EA) were also factored out. Finally, we included unmodulated predictors to capture any unaccounted variance in the BOLD signal, especially from areas associated with early encoding of the stimulus evidence itself (Fig. 6a).

**Figure 6.**
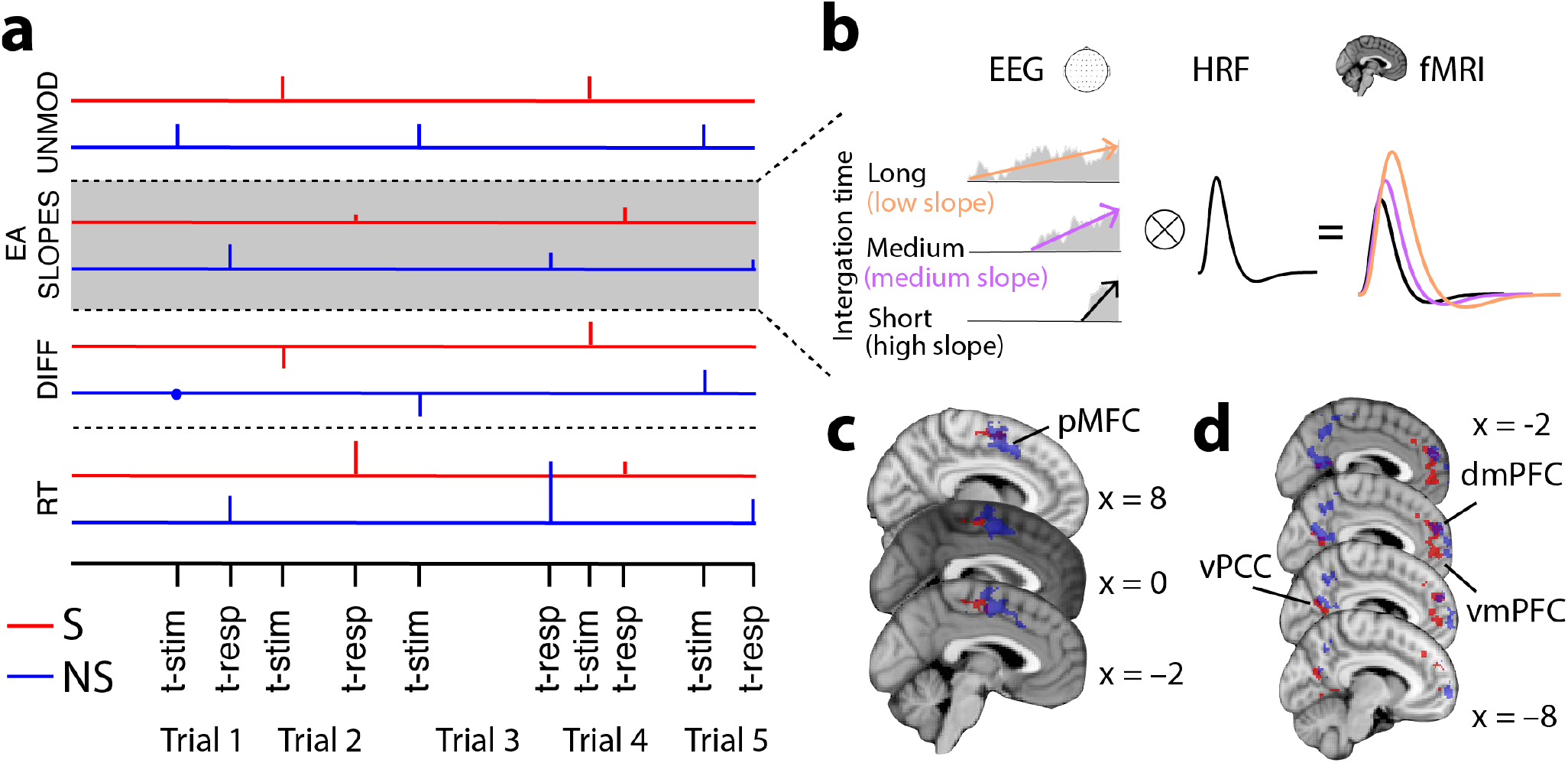
EEG-informed fMRI analysis. **a**. The fMRI GLM model included two parametric boxcar regressors capturing the electrophysiological trial-by-trial variability in the slope of evidence accumulation (EA) for each of the social and non-social trials at the time of decision. To absorb the variance associated with other task-related processes – for each of the social and non-social trials separately – we included the following additional regressors: UNMOD – two unmodulated (amplitude 1) boxcar regressors at the onset of the stimuli, DIFF – two parametric boxcar regressors of task difficulty (−1: hard, 0: mid, 1: easy), and RT – two parametric boxcar regressors modulated by individual trial RTs at the time of decision. All regressors had a fixed duration of 100 ms. **b**. Example EA traces with different build up rates (coloured arrows). Convolving these traces with a hemodynamic response function (HRF) leads to higher predicted fMRI activity for longer compared to shorter integration times – that is, higher predicted fMRI activity for shallower compared to steeper EA slopes. **c**. According to the hypothesized prediction in textbfb) the EEG-informed fMRI predictors of the slope of EA revealed an activation in posterior medial-frontal cortex (pMFC) for both social and non-social trials. **d**. We found that the pMFC showed task-dependant co-activation with regions of the human valuation system, specifically clusters along the medial wall of the prefrontal cortex as well as regions of the posterior cingualte cortex. The clusters represent mixed-effects activations that survived |*Z*| > 2.57 and that were cluster-corrected (*P* < 0.05) using a resampling procedure with a minimum cluster size of 88 voxels (see Materials and Methods). The complete lists of activations are shown in Supplementary Tables 1 and 4. vmPFC: ventromedial profrontal cortex; dmPFC: dorsomedial prefrontal cortex; vPCC: ventral posterior cingualte cortext.

Our EEG-informed fMRI approach benefits from using the actual neural signals, which could capture latent variability in information processing that might otherwise go amiss when using simple behavioral or model-derived indices (Sajda, Philiastides, Heekeren, and Ratcliff, 2011). For example, most sequential sampling models only produce mean estimates of the relevant decision variables (e.g. drift rate) across many trials, with only few studies attempting to derive single-trial parameter estimates (Gluth, Hotaling, and Rieskamp, 2017; Turner, Van Maanen, and Forstmann, 2015). Here, instead, we estimate the rate of EA on individual trials purely from the slope of the accumulating activity we identified in the EEG data. This way we could account for true endogenous variability in EA and circumvent potential issues related to model estimation and/or mis-specification when deriving build-up rates purely based on fits to behavior.

Crucially, trials with lower EA rates that require longer integration times to reach the decision boundary should have larger areas (energy) under the accumulation curve (Basten, Biele, Heekeren, and Fiebach, 2010; Hare et al., 2011; Liu and Pleskac, 2011). Correspondingly, we hypothesize that candidate accumulator regions should appear to be more hemodynamically active in trials with longer compared to shorter integration times (Fig. 6b). This, in turn, should be consistent with a negative relationship between our EEG-informed EA slope predictor and the BOLD response in the relevant brain areas (Hare et al., 2011; Liu and Pleskac, 2011; Mulder, Van Maanen, and Forstmann, 2014). Consistent with a domain-general EA neural architecture, we identified a region of the posterio-medial frontal cortex (pMFC) correlating negatively with the trial-by-trial EA slope in our EEG-informed predictor (Fig. 6c) in both both social and non-social contexts. We note that in non-social choices the cluster extended more anterior compared to social choices, suggestive of a potential gradient organisation within the pMFC associated with EA. We found no domain-specific activations surviving in the direct contrast between social and non-social contexts for the EA slope predictor.

To further establish the role of the pMFC in EA across social and non-social contexts, we aimed to demonstrate whether its activity exhibits a task-dependent coupling with brain regions encoding the relevant decision evidence and the extent to which this coupling arises from domain-general or domain-specific neural representations. To this end, we ran separate psychophysiological interaction (PPI) analyses for each of the social and non-social trials, using the context-specific pMFC clusters as seed and the trial-wise task difficulty as the psychological predictor (see Materials and Methods for more details). We hypothesize that the relevant coupling with pMFC should be negative, as easier trials decrease integration times and correspondingly the overall integrated activity (that is, area under the accumulation curve; Fig. 6b).

The PPI analyses from both social and non-social contexts revealed significant negative coupling (by task difficulty during the decision phase) between the pMFC and regions of the human valuation system. More specifically, we observed activations in posterior cingulate cortex (PCC) as well as in dorso- and ventro-medial prefrontal cortex (dmPFC/vmPFC; Fig. 7), consistent with recent resting-state connectivity reports showing negative BOLD correlations between these regions and the pMFC (Neubert, Mars, Sallet, and Rushworth, 2015). Intriguingly, these areas have repeatedly been implicated in encoding a “common neural currency” of abstract value signals used in the process of EA (Pearson, Watson, and Platt, 2014; Piva et al., 2019; Rangel and Hare, 2010). As with the EA clusters, we found that in the non-social context, activations were situated more anterior relative to the social context, consistent with previous reports of value gradients within the medial prefrontal cortex (Chib, Rangel, Shimojo, and O’Doherty, 2009; Clithero and Rangel, 2014; D. V. Smith et al., 2010). Taken together, our findings provide compelling evidence that relevant decision evidence is converted into a “common currency” along the medial wall of the human brain and subsequently integrated for the decision in the pMFC. We found no domain-specific activations surviving in the direct contrast between social and non-social contexts for the PPI predictor.

**Figure 7.**
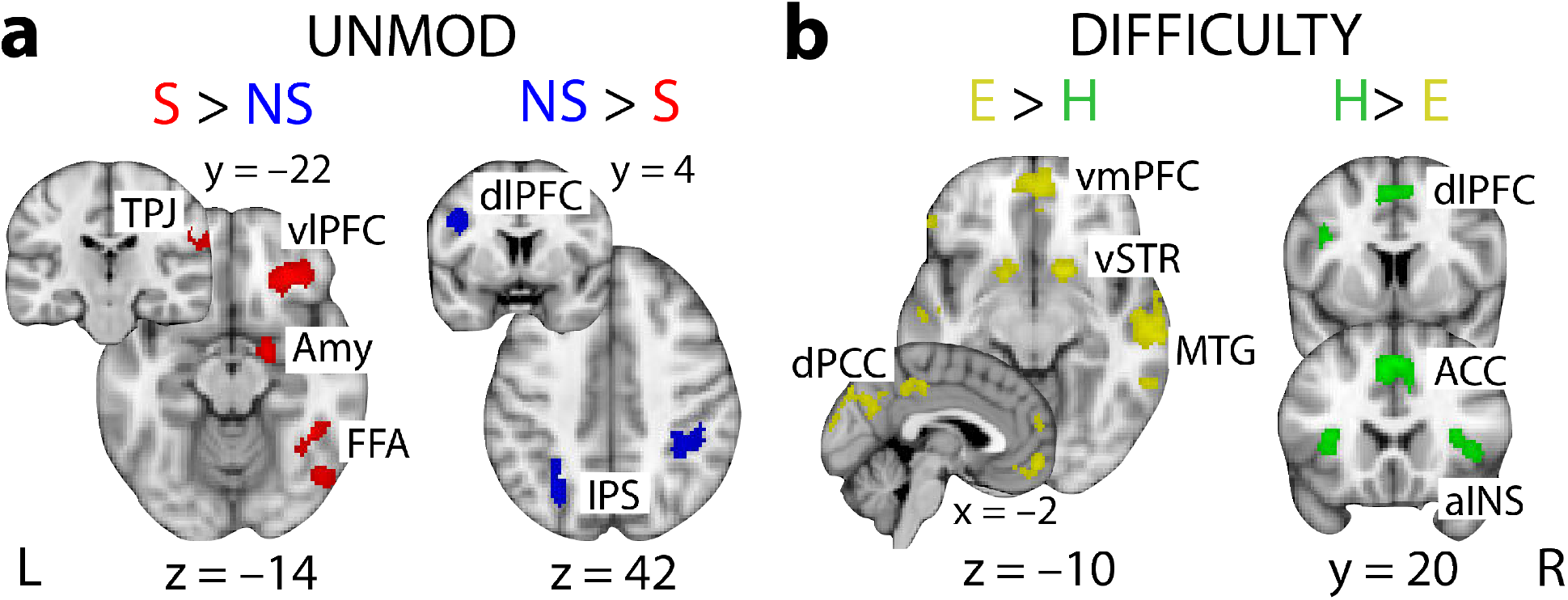
Domain-specific and task difficulty fMRI activations. **a**. Activations showing greater BOLD response for social than non-social trials (red) and those exhibiting higher response for non-social compared to social trials (blue). These activations arise from the contrast of the two unmodulated regressors (UNMOD) in Fig. 6a. **b**. Activations showing greater BOLD response for easy than difficult trials (yellow) and those exhibiting higher response for difficult compared to easy trials (green). These activations arise from the conjunction of the two task difficulty regressors (DIFF) for social and non-social trials in Fig. 6a. All clusters represent mixed-effects activations that survived |*Z*| > 2.57 and that were cluster-corrected (*P* < 0.05) using a resampling procedure with a minimum cluster size of 88 voxels (see Materials and Methods). The complete list of activations is shown in Supplementary Tables 2 – 3. vlPFC: ventrolateral prefrontal cortex; Amy: amygdala; FFA: fusiform face area; TPJ: temporoparietal junction; IPS: intraparietal sulcus; vmPFC: ventromedial prefrontal cortex; vSTR: ventral striatum; MTG: medial temporal gyrus; dPCC: dorsal posterior cingulate cortex; dlPFC: dorsolateral prefrontal cortex; ACC: anterior cingulate cortex; aINS: anterior insula.

#### Domain-specific and task difficulty neural representations

To identify brain areas processing domain-specific representations (e.g. areas encoding the initial evidence for each of the two contexts prior to conversion to a “common currency”) we examined the contrast between the unmodulated predictors for the social vs. non-social trials (Fig. 6a). We identified a distributed set of regions activating stronger for social compared to non-social trials, in the right fusiform gyrus, right amygdala and right ventrolateral prefrontal cortex (collectively referred to as the “face network” Skelly and Decety, 2012) as well as in the right temporoparietal junction (TPJ) (Fig. 7a). These findings are consistent with the processing of facial characteristics required for mentalizing and inferring the opponent’s intentions in the social trials, respectively (Cerniglia et al., 2019). In contrast, non-social trials exhibited increased activation patterns relative to non-social trials in areas of the lateral intraparietal cortex bilaterally as well as in the left dorsolateral preforntal cortex (dlPFC) (Fig. 7a), which have both been implicated in encoding risk and reward probabilities in non-social contexts (Burke and Tobler, 2011; Daw et al., 2006; B. W. Smith et al., 2009).

Task difficulty in our task was proportional to the reward probability associated with a ’Play’ choice. We, therefore, expected the task difficulty predictor to correlate both positively (i.e. easy > difficult) with areas known to encode the value of a given choice and negatively (i.e. difficult > easy) with regions of the human attentional network encoding overall task demands. Across both social and non-social contexts, we found positive correlations with regions of the human valuation system such as the ventromedial prefrontal cortex (vmPFC), ventral striatum and the posterior cingulate cortex (Clithero and Rangel, 2014; Domenech, Redouté, Koechlin, and Dreher, 2018) (Fig. 7b). Consistent with previous reports (Grinband et al., 2008; Monosov, 2017; Philiastides and Sajda, 2007), we also found negative correlations with regions encoding uncertainty and attentional control such as the anterior cingulate cortex, lateral prefrontal cortex and anterior insula in both social and non-social contexts (Fig. 7b). We note, that though these results are consistent with a long body of previous work, they are nonetheless important to report here as further validation of the choice of our fMRI analysis design, which offered a reliable account of all relevant experimental manipulations.

## Discussion

The marriage of social and non-social forms of uncertainty into a comprehensive theory of decision-making promises to significantly improve our understanding of human behavior. To date, however, there is little work directly comparing the two forms of uncertainty during economic decision-making. Correspondingly, the extent to which social and non-social forms of uncertainty drive decisions via a common mechanism and network architecture remains elusive. As a result, two competing theoretical frameworks have recently been proposed to help identify differences and similarities across the neural processes that guide social and non-social choices (Ruff and Fehr, 2014).

The “common currency” framework posits that a common neural architecture encodes an integrated value of all factors guiding each of the social and non-social choices, although the perceptual information that leads to these unified value representations might be represented in domain-specific brain regions. In contrast, the “social valuation” framework proposes that value information associated with social and non-social choices is processed and integrated in dedicated neural circuits, albeit with similar computational principles, in line with the social brain hypothesis (Dunbar and Shultz, 2007).

Our work offers critical new insights that could help arbitrate between these two accounts. More specifically, our results are more consistent with the “common currency” framework, whereby domain-specific perceptual information associated with each domain is likely converted into a common neural currency in regions of the human valuation system before being integrated for the decision in medial frontal cortex. A key feature of our work – enabling fair and direct comparisons between social and non-social choices – was the embedding of both choice types in the context of an economic game in which task difficulty varied along the same scale of reward probability across social and non-social stimuli.

Behavioral performance on the task as well as modelling of the choice-RT data using a simple sequential sampling model were suggestive of comparable decision dynamics across social and non-social contexts. Correspondingly, we identified ramp-like activity in the electrophysiological signal with a well-demarcated centroparietal scalp profile that were virtually identical across the two contexts. We further demonstrated that trial-by-trial variability in the slope of this accumulating activity was highly predictive of choice behavior. These findings point to a domain-general accumulation-to-bound decision mechanism (Bogacz et al., 2006), consistent with previous reports in value-based (Gluth, Rieskamp, and Büchel, 2012; Pisauro, Fouragnan, Retzler, and Philiastides, 2017; Polaniía, Krajbich, Grueschow, and Ruff, 2014), perceptual (Drugowitsch et al., 2012; Gherman and Philiastides, 2015; Kelly and O’Connell, 2013) and even memory-based decisions (van Vugt, Beulen, and Taatgen, 2019).

Another novel aspect of our work is that we exploited the endogenous variability in the process of evidence accumulation to identify candidate regions involved in this process. More specifically, we used the trial-wise changes in the rate of evidence accumulation – which embody the momentary changes in the decision process as it unfolds – to predict BOLD activity in the fMRI signal. Note that, unlike conventional event-related potentials for which the trial-by-trial signal fluctuations might be contaminated from unspecific neural process, the nature of our multivariate discriminant analysis ensured these processes were effectively subtracted out, facilitating instead the spatial “unmixing” of the main signal of interest (i.e. process of evidence accumulation) (Philiastides, Heekeren, and Sajda, 2014; Ratcliff, Philiastides, and Sajda, 2009).

Critically, our approach differs from conventional fMRI analyses in that it capitalizes on the trial-by-trial fluctuations in electrophysiologically-derived temporal components that might not be fully reflected in behavior. In turn, this approach could offer additional explanatory power than what can be achieved by purely stimulus- or behaviorally-derived predictors. In deploying this EEG-informed fMRI analysis, we were able to identify a region of the pMFC correlating robustly with the trial-by-trial changes in the rate of evidence accumulation across both social and non-social contexts, suggesting that both types of decisions are likely to rely on the same implementational and algorithmic process (Lockwood et al., 2017).

The pMFC cluster we identified here lies on the medial surface of the juxtapositional lobule cortex in a region commonly referred to as the supplementary motor area and extending into portions of the adjacent pre-supplementary motor area, bilaterally. Both areas have traditionally been linked to preparing voluntary actions (Kim et al., 2010; Nachev, Kennard, and Husain, 2008) but have also been assigned other functional roles, including decision boundary adjustments in the context of accumulation-to-bound models (Forstmann et al., 2008; Ivanoff, Branning, and Marois, 2008). Our results suggest that the role of the pMFC extends beyond mere decision boundary adjustments and involves instead the encoding of the full temporal dynamics of the process of evidence accumulation, a process that might be mediated by an increased tendency to select the appropriate motor response.

Correspondingly, our results support the rapidly emerging view that, at least under conditions of increased urgency to make a choice, decisions are embodied in the same sensorimotor areas guiding the actions used to express that choice. This interpretation is consistent with past neuroimage findings (Donner, Siegel, Fries, and Engel, 2009; Filimon et al., 2013) and recent computational accounts ascribing an active “motor accumulation” role to (pre)motor structures (Steinemann, O’Connell, and Kelly, 2018; Verdonck, Loossens, and Philiastides, 2021). Similarly, recent animal experiments have shown that while neural representations in sensory and association cortices are sufficient to perform a simple discrimination task, inactivation of downstream (pre)motor areas can led to gross behavioral impairments (Peixoto et al., 2021; Z. Wu et al., 2020).

To further probe the implementational level of social and non-social decisions, we performed a psycho-physiological interaction (PPI) analysis. This analysis allowed us to investigate the functional coupling of the pMFC with other brain regions to provide further insights that would help arbitrate between the “common currency” and “social valuation” accounts introduced above. We found that pMFC activity exhibited task-dependent coupling (i.e. as the process of evidence accumulation unfolds) with regions of the human valuation system in the dmPFC, vmPFC and PCC. This coupling was present in both social and non-social choices, consistent with the relevant decision evidence being converted into a “common currency” prior to being used in the process of evidence accumulation to drive the final commitment to choice.

While pMFC activity covaried systematically with areas of the human reward network, we nonetheless observed some degree of spatial dissociation across contexts along an anterior/posterior axis within these areas, consistent with recent reports from human and animal work advocating for a gradient-based organization along the medial wall of the brain (Kolling et al., 2021). Specifically, in the social context the relevant clusters were situated within the most posterior sections of the medial prefrontal cortex (Ferrari et al., 2016; Lieberman et al., 2019) whereas those of the non-social context occupied sections of the most anterior portions of this region (Chib, Rangel, Shimojo, and O’Doherty, 2009; Clithero and Rangel, 2014; D. V. Smith et al., 2010). Interestingly, this organization seems to be preserved along the cascade of constituent processes, from the relevant value representation to the process of evidence accumulation.

The common currency account also postulates that prior to embedding within a domain-general valuation system, social and non-social choices might give rise to domain-specific early representations (Hutcherson, Montaser-Kouhsari, Woodward, and Rangel, 2015; Lim, O’Doherty, and Rangel, 2013). Here, we observed that social choices were associated with increased activity in the FFA, the amygdala and vlPFC – with a right lateralization – consistent with the well-known “face network”, which is crucial for face identification and affective processing of faces (Garvert, Friston, Dolan, and Garrido, 2014; Vuilleumier et al., 2004). Similarly, we observed increased activation in the right TPJ, a region implicated in social cognition and various types of mentalizing relevant for extracting semantic meaning (value) from an opponent’s face identity (Van Overwalle, 2009). Conversely, the non-social condition was uniquely linked to activity in the lateral intraparietal cortex which has been linked to the encoding of pure reward probabilities both in primates (Burke and Tobler, 2011; Sugrue, Corrado, and Newsome, 2004) and humans (Daw et al., 2006; S.-W. Wu, Delgado, and Maloney, 2015), suggesting that these choices were guided by the consideration of the likelihood of receiving a reward as presented explicitly during the task.

In conclusion, our results offer compelling new evidence that social and non-social choices share common neural underpinnings, whereupon domain-specific information is converted into a “common currency” in domain-general valuation areas prior to being accumulated for the decision in medial frontal cortex. Similarly, our multimodal research approach – including the fusion of EEG and fMRI – offers news opportunities for a more targeted characterization of the computational principles and the neural systems involved in human decision making under different forms of uncertainty. As optimal decision making is at the heart of strategic planning, providing a mechanistic account of decision making under risk and uncertainty could have wider, long-term, socioeconomic impact and facilitate an improved understanding of maladaptive choice behavior.

## Materials and Methods

### Participants

40 participants were recruited through the The University of Glasgow subject pool. Since facial perception may depend on one’s race and racial history (e.g. Scott and Monesson, 2009), participants were chosen to be Caucasians, aged 18-35 to match the available face stimuli (see below). Two participants were removed due to the poor behaviour (one had near chance performance across all levels of reward probability in the social context whereas the other had chosen to nearly always ’Play’ across all levels of reward probability in the non-social context) and seven participants due to noisy EEG signals in the scanner leading to poor (chance) discrimination performance. The remaining 31 subjects (12 males, 19 females), were included in all subsequent analyses. They all had normal or corrected-to-normal vision and reported no history of psychiatric, neurological or major medical problems, and were free of psychoactive medications at the time of the study. The study was approved by the College of Science and Engineering Ethics Committee at the University of Glasgow (300180147) and informed consent was obtained from all participants.

### Stimuli

We used a set of 150 photorealistic face images (400 × 300 pixels). We presented all stimuli centrally via an LCD projector (frame rate = 60 Hz) on a screen placed at the rear opening of the bore of the MRI scanner, and viewed through a mirror mounted on the head coil (distance to screen = 95 cm), using Presentation software (Neurobehavioral Systems Inc., Albany, CA). These face images were assigned to the five levels of reward probabilities used in the main task, based on participant-specific indirect trustworthiness ratings for each face (see Procedure below). Nonetheless, to encourage a broad range of indirect trustworthiness reports from our participants, we manipulated these images to obtain versions across a potentially wide range of trustworthiness levels.

Specifically, we used a reverse correlation procedure (Ahumada Jr and Lovell, 1971) to first identify features associated with higher trustworthiness scores and we then manipulated these features in all faces to create different trustworthiness versions of each face using a Generative Face Grammar (Yu, Garrod, and Schyns, 2012). We estimated the facial features associated with trustworthiness judgements from a separate set of 416 faces that were each first 3D-captured using a Di4D (Dimensional Imaging, Glasgow, UK) facial capture system and subsequently rendered and rated for their trustworthiness by 49 independent observers, based on a reverse-correlation procedure reported in (Zhan, Garrod, van Rijsbergen, and Schyns, 2019).

Each captured face is described by a (4735*3 × 1) vector of 3D mesh vertex coordinates (x,y,z) and a (800*600*3 × 1) vector of texture pixel colours (r,g,b). For each mesh vertex coordinate and texture pixel we fit a multiple linear regression model predicting the value of the coordinate or pixel as a function of the following predictors: average trustworthiness rating over the 49 observers, age of the face, sex of the face, ethnicity of the face, plus a constant term and interaction terms between the predictors, yielding a set of linear coefficients for each vertex coordinate and pixel. Categorical predictors were coded as one-hot vectors, continuous predictors were coded as their real values. Each individual face vertex and texture pixel is then described by the output of this linear model at the given values of the predictors plus an identity-specific residual term.

To generate individual faces of varying degrees of perceived trustworthiness for a specific identity we evaluated the linear model at varying levels of the trustworthiness predictor while holding the remaining predictors constant (at the observed values for this identity) and finally add the identity-specific residual term and render the resulting 3D model. This procedure was first applied to 131 identities, made up of 61 male images, 70 female images (all Caucasian), and an additional set of 19 new identities were included (to increase the image sample size to 150) by perturbing the identity-specific residuals in the above procedure with noise in the directions of the principal components of the identity space.

Using this procedure we created twenty trustworthiness versions for each face, 1 representing the least trustworthy version and 20 displaying a trustworthy version of the face. Only one version per face identity (original and fabricated) was chosen for the main experiment. As noted above, this procedure was adopted purely for increasing the likelihood that stimuli would fall into a wide range of different trustworthiness categories, though ultimately the final categorization of each face was based on the participant-specific reports as we explain in more detail below.

### Experimental paradigm

The experimental design consisted of three parts: 1) an initial behavioral session comprising a rating task and a separate choice task, 2) an online rating task one day prior to the main EEG-fMRI experiment and 3) a rating task and the main choice task during which participants underwent scanning. All tasks were framed in the context of an economic game. Specifically, we used a variant of the trust game in which participants engaged in a series of one-shot trust games involving a Trustee and an Investor. In each game, the Trustee is allocated 1 point per trial and has two options: 1) to obtain a small but certain reward by keeping the 1 point (’Keep’ option) or 2) to invest the 1 point with the Investor for a bigger but uncertain reward (’Play’ option). When the Trustee chooses the latter option the 1 point is quadrupled and it is now up to the Investor to determine whether to keep all 4 points for themselves or split them evenly between the two players 2.

During the rating tasks, we told participants that the face identities belonged to individuals who have previously taken part in an economic game (i.e. a trust game like the one described above) in which they were assigned the role of Investor. The goal of the participants in these indirect trustworthiness rating tasks was to assess the face identities’ social attitudes by estimating the overall likelihood with which each Investor split the augmented endowment (in the range of 0-1, on a continuous scale). This framing ensured that our social stimuli varied along the same scale of reward probability as non-social gambles (see below). This was a critical feature of our design since embedding the rating in the context of a trust game allowed us to circumvent the arbitrary nature of explicit trustworthiness ratings (e.g. using likert scales) and ensured a direct mapping between social and non-social choices. Importantly, when instructing participants, we purposely avoided explicit mentions of “trustworthiness” to sidestep the possibility of participants developing unusual strategies in the game due to social desirability biases.

Correspondingly, we told participants that during the main choice task they would assume the role of the Trustee themselves and play with the same face identities they encountered during the ratings tasks (in social trials) or using purely probabilistic gambles (in non-social trials). On each trial participants had to choose between the ’Keep’ and ’Play’ option, however the outcome of the trial depended on the context. In the social context, we told participants that the probability of doubling their points was based on one of the Investor’s responses from when they previously played the game in our lab. In reality, we used the participant-specific reports on the likelihood of individual face identities splitting the augmented endowment to construct reward probability ranges that were comparable to those used in the non-social contexts (via explicit reward probability values). This design ultimately ensured that participants’ decisions in the main choice task would be based on the same economic considerations – that is, the reward probability associated with a ’Play’ choice – across both contexts. To make our cover story more realistic for our participants, we took pictures of their own faces and told them that their face displays and responses would be used for similar experiments in the future. In reality, we would delete the pictures after each session.

In the main choice task we included five different levels of reward probabilities (given a ’Play’ choice); 0–0.2, 0.2–0.4, 0.4-0.6, 0.6–0.8 and 0.8–1. In the social context, we assigned each of the face identities into the five bins based on the participants ratings on the day of the EEG-fMRI. We nonetheless used their ratings across all three ratings sessions to identify face identities that received inconsistent ratings across sessions (more than two bins apart) and removed them from the experiment (on average, 10.807 face identities were removed). On average, there were were 23, 34, 32, 36 and 14 face identities across the five levels of reward probability, respectively. In the non-social context, the reward probability was presented explicitly through a probability range displayed on a face neutral for trustworthiness. We embedded a background face in the non-social trials to equalize the perceptual load across contexts to enable direct comparisons between contexts (e.g. when searching for domain-specific representations). We additionally distorted the number presenting the reward probability ranges to equalize the early encoding of the perceptual stimuli across contexts (by ensuring that RTs and non-decision times were comparable across contexts, during initial piloting of the task). During these trials, we instructed participant to focus exclusively and base their choices on the numbers displayed on the stimuli.

The main choice task during the initial behavioral session was virtually identical to the one used for the EEG-fMRI experiments, with the main difference being that participants received feedback following each of their choice on a trial-by-trial basis. We included this initial choice task in order to identify participants who would understand the task and to offer participants a chance to familiarize themselves with the main paradigm. We provided immediate feedback relating to their choice to reinforce the associations between the stimuli and the outcome. During the EEG-fMRI experiments participants would be informed of their accumulated points only during the breaks (after every 50 trials). Furthermore, to motivate participants to engage with the task, we told them that in addition to their base rate payment (behavioral session: £6, EEG-fMRI session: £16) they would receive a variable bonus (up to £4) based on the overall points they collected during the experiment. We did not provided further details on how points translated into monetary rewards. On average, the participants who were included in the final analysis received £8.03 ± 0.31 during their initial behavioral session. In the EEG-fMRI session, we paid all participants £20.

The main choice task during the EEG-fMRI session comprised of a total of 500 trials, split evenly across the social and non-social contexts. All trials were presented in fully-interleaved fashion and they were further broken down into 5 runs, lasting approximately 7 minutes each. In each run we included a 30 second break at the middle and the end (i.e. after every 50 trials). During each trial, a jittered fixation cross (1-4s, mean = 2s; optimised for maximizing discriminability across contexts as in (Mumford, Poline, and Poldrack, 2015) would be followed by the presentation of the face stimulus (either social or non-social). For both contexts, the stimulus remained on screen until the participant made a response or for a maximum of 1.3 seconds. If the participant’s response was faster than 1.3s, the fixation cross reappeared for the remaining time up to 1.3s, in order to keep the run times consistent between participants. Participants used an MR-compatible response box (Cambridge Research Systems, 2019) to indicate their choices.

### Choice probabilities

To assess the similarity between the probabilities of ’Play’ choices across the social and non-social contexts we used a conventional likelihood-ratio test. Specifically, we examined whether a single sigmoid curve (Weibull function) would fit the combined social and non-social choice data across the five reward probability levels better than two separate curves (Philiastides and Sajda, 2006). We performed this separately for each participant by fitting the best single Weibull function jointly to the two data sets in addition to the individual fits. The likelihoods (L) obtained from this procedure were transformed using the following equation:

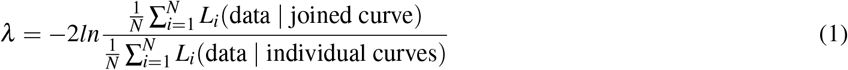

where *N* represents the number of participants and *λ* is distributed as *χ*^2^ with two degrees of freedom (Hoel, Port, and Stone, 1971). If *λ* exceeds the criterion value (for *p* = 0.05), we concluded that a single function fits the data better than two separate domain-specific functions.

### Sequential sampling modelling

We modelled the behavioral data using a special case of the leaky competing accumulator model. Specifically, we used an Ornstein–Uhlenbeck process to model the evidence accumulation (EA) stage as in (Pisauro, Fouragnan, Retzler, and Philiastides, 2017; Polaniía, Krajbich, Grueschow, and Ruff, 2014):

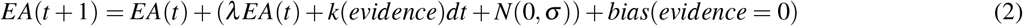

We set the decision thresholds for ’Play’ and ’Keep’ choices to +1 and −1 respectively such that positive drift rates are associated with reward probability levels favoring ’Play’ choices while negative drift rate values with those favoring ’Keep’ choices. Correspondingly, in Eq. 2 the evidence represents a transformed version of the original five reward probability levels such that they are now centered around zero (i.e. [−0.5 −0.25 0 0.25 0.5]).

Free parameter *k* modulates the input, *λ* represents the acceleration to threshold and *N*(0, *σ*) is a Gaussian noise term with standard deviation *σ*. We used a time increment dt = 0.001s and assumed that the model makes a decision when |EA| >boundary. Early visual encoding and motor preparation were captured by the non-decision time free parameter (nDT), which was included into the total reaction time (RT). For the indecision point (i.e. 0 evidence) we included an extra free parameter, *bias*, to account for potential inter-individual biases towards either ’Play’ or ’Keep’ choices. We separated the RTs according to the selected action (’Keep’ or ’Play’). We then combined the RTs from both trial types into a single distribution by flipping the sign of the ’Keep’ trials, so that all the times in this distribution received a negative sign (Voss, Rothermund, and Voss, 2004). This RT distribution and participants’ choice probabilities were compared to the RT distribution and proportion ’Play’ choices generated by the model. For a given set of parameter estimates, we estimated the log likelihood (LL) of the data using the following formula:

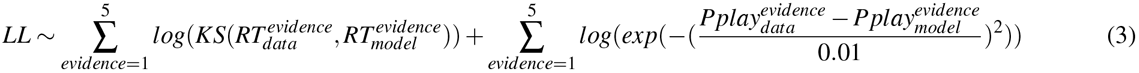

KS(p,q) is used to estimate the probability that two distributions are equal, based on the Kolmogorov–Smirnov test (via the ktest2 function in MATLAB). Pplay represents the fraction of ’Play’ choices for each of the five levels of evidence. To fit the model we used a two step procedure. Firstly, we used the fmincon MATLAB function to provide an initial estimate of participant-specific parameters. Specifically, we ran this procedure 20 times and the parameters associated with the smallest LL were selected for the next step. secondly, we ran a grid search fitting procedure for each participant using a fine-grained parameter space around the estimates obtained in the previous step. Choices and RT distributions were created for each possible combination of the four free parameters from 5000 simulated decision traces for context.

Mean parameter estimates for the social context: lambda: 5.774 ± 2.357, k: 3.206 ± 1.555, sigma: 0.02 ± 0.01, bias: −0.00004 ± 0.0004, nDT: 0.336 ± 0.09. Mean parameter estimates for non-social context: lambda: 5.277 ± 2.37, k: 2.611 ± 1.355, sigma: 0.011 ± 0.006, bias: −0.00002 ± 0.0006, nDT; 0.304 ± 0.089. For the majority of parameters, there was no significant differences between the the two contexts (*λ* : t(30) = −1.3, p = 0.203, bias: t(30) =0.26, p = 0.8, k: t(30) =-1.349, p = 0.188, nDT: t(30) =-1.363, p = 0.183), reinforcing the notion that choice behavior was comparable across social and non-social choices. There was a small but significant difference in the noise term (σ: t(30) =-4.244, p<0.001), likely due to additional internal variability in processing the faces and their trustworthiness in the social context compared to processing the numbers in the non-social trials.

### EEG data acquisition

We used an MR-compatible EEG amplifier system (Brain Products, Germany) to collect the data and Brain Vision Recorder software (Brain Products, Germany) to continuously record EEG at 5000 Hz. A hardware 0.016-250Hz band-pass filtered the data online. We placed the 64 Ag/AgCl scalp electrodes according to the 10 – 20 system, with the reference and ground electrodes being built in between electrodes Fpz and Fz and between electrodes Pz and Oz, respectively. Each electrode had in-line 10 kOhm surface-mount resistors to ensure subject safety, which was further guaranteed by bundling and twisting all leads for their entire length. We lowered the input impedance for each electrode to < 50 kOhm (25 KOhm average across participants). The acquisition of EEG and MRI data was synchronized (Syncbox, Brain Products, Germany) and MR-scanner triggers were recorded separately for the subsequent offline removal of MR gradient artifacts. To facilitate the recording of the scanner triggers, the scanner pulses were lengthened to 50 *μ* s via an in-house pulse stretcher. Experimental event codes and participants’ responses were synchronized, and recorded simultaneously, with the EEG data through the Brain Vision Recorder software. We positioned subjects inside the scanner by ensuring that electrodes Fp1 and Fp2 were aligned with the isocentre of the MR scanner. Finally, the cabling connecting to the EEG amplifiers at the back of the bore was secured to a cantilever beam to minimize scanner vibration artifacts.

### EEG data preprocessing

We used MATLAB (Mathworks, Natick, MA) to preprocess and analyse the EEG data. EEG signals recorded inside an MR scanner are contaminated primarily with MR gradient artifacts and ballistocardiogram (BCG) artifacts due to magnetic induction on the EEG leads. To correct for gradient-related artifacts, we constructed average artifact templates from sets of 70 consecutive functional volumes centred on each volume of interest, and subtracted these from the EEG signal. This process was repeated for each functional volume in our dataset. We removed any residual spike artifacts we applied a 12 ms median filter. Furthermore, we applied a 0.5 – 20 Hz band-pass filter in order to remove slow DC drifts and higher frequency noise. All data were downsampled to 1000 Hz.

To remove eye blinks we asked participants to perform an eye movement calibration task prior to the main experiment during which they were instructed to blink repeatedly several times while a central fixation cross was displayed in the centre of the computer screen. We recorded the timing of these events and used principal component analysis to identify linear components associated with eye-blinks, which were subsequently removed from the broadband EEG data collected during the main task (Parra, Spence, Gerson, and Sajda, 2005).

BCG artifacts share frequency content with the EEG and are therefore more challenging to remove. To avoid loss of signal power in the EEG we only removed a small number of participant-specific BCG components using principal component analysis and relied instead on our single-trial classifiers (see single-trial EEG analysis section below) to identify task-related discriminating components that are likely to be orthogonal to the BCG (Fouragnan, Retzler, Mullinger, and Philiastides, 2015; Gherman and Philiastides, 2018). This approach is robust to the presence of BCG artifact residuals, specifically, due to the (spatial) multivariate nature of our classification techniques.

Correspondingly, we first extracted BCG principal components from the data after low-pass filtering at 4Hz (i.e. to extract the signal within the frequency range where BCG artifacts are typically observed) and then created datasets with different number of principal components removed (up to 5). The sensor weightings corresponding to the relevant components were projected onto the broadband data and subtracted out. We determined the number of optimal principal components for each participant by maximizing classification performance along the task-relevant dimension (see below) using cross validation (average number of BCG components across participants: 2.447 ± 1.969).

### Single-trial EEG analysis

To identify activity related to evidence accumulation we used a single-trial multivariate discriminant analysis (Parra, Spence, Gerson, and Sajda, 2005; Sajda, Philiastides, and Parra, 2009) to discriminate between easy (i.e. reward probabilities 0–0.2 and 0.8–1) and difficult trials (reward probabilities 0.4–0.6) in stimulus-locked EEG data, collapsing across both social and non-social trials. We were particularly interested in identifying accumulating activity with a build-up rate proportional to the amount of decision difficulty that would have in turn enabled a gradual increase in the discriminator’s performance while the traces for easy and difficult trials diverged as a function of elapsed time in stimulus-locked data (Fig. 4a). We treated the medium difficulty trials (i.e. reward probabilities 0.2–0.4 and 0.6–0.8) as “unseen” data, to more convincingly test for a full parametric effect on the build-up rate associated with the different levels of decision difficulty (see below).

More specifically, our method estimates an optimal combination of EEG sensor linear weights (i.e., a spatial filter **w**) which, applied to the multichannel EEG data (**x(t)**), yields a one-dimensional projection (i.e., a discriminant component *y*(*t*)) that maximally discriminates between the two contexts:

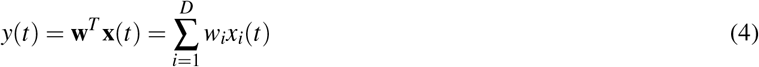

where *D* represents the number of channels, indexed by *i*, and *T* indicates the transpose of the matrix. We applied this method to identify **w** for short (60 ms) overlapping time windows centred at 10 ms-interval time points, between −100 and 800 ms relative to the onset of the decision stimulus. This procedure was repeated for each subject and time window separately. By integrating information spatially across the multidimensional sensor space, we increase signal-to-noise ratio whilst simultaneously preserving the trial-by-trial variability in the relevant discriminating component. More specifically, applied to an individual trial, spatial filters (**w**’s) obtained in this way produce a measurement of the discriminant component amplitude for that trial, which we treat as a neural surrogate of the relevant decision activity.

To estimate the optimal discriminating spatial weighting vector **w** we used a regularized Fisher discriminant analysis as follows: **w** = **S*_c_***(**m**_2_ – **m**_1_), where *m_i_* is the estimated mean of condition i and *S_c_* = 1/2(*S*_1_ + *S*_2_) is the estimated common covariance matrix (i.e., the average of the condition-wise empirical covariance matrices, 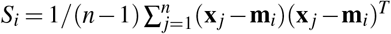, with *n* = number of trials). We replaced the condition-wise covariance matrices with regularized versions of these matrices to counteract potential estimations errors: 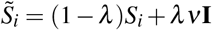, with *λ* ∈ [0, 1] being the regularization term and *v* the average eigenvalue of the original *S_i_* (i.e., *trace*(*S_i_*)/*D*, with *D* corresponding to the dimensionality of our EEG space). Note that *λ* =0 yields unregularized estimation and *λ* = 1 assumes spherical covariance matrices. Here, we optimized *λ* for each participant using leave-one-trial-out cross validation with the following *λ* values ∈ [0, 0.01, 0.02, 0.04, 0.08, 0.16] (*λ* mean ± s.e.: 0.067 ± 0.072).

To quantify the performance of the discriminator for each time window, we computed the area under a receiver operating characteristic (ROC) curve (i.e., the *A_z_* value), using a leave-one-trial-out cross-validation procedure. Specifically, for every iteration, we used N-1 trials to estimate a spatial filter (**w**), which we then applied to the remaining trials to obtain out-of-sample discriminant component amplitudes (*y*(*t*)). We used these out-of-sample amplitudes to compute the *A_z_*. In addition, we determined participant-specific *A_z_* significance thresholds (rather than assuming an *A_z_* = 0.5 as chance performance) using a subsequent bootstrap analysis whereby trial labels were randomised and submitted to the leave-one-trial-out test described above. This randomisation procedure was repeated 500 times, producing a probability distribution for *A_z_*, which we used as reference to estimate the *A_z_* value leading to a significance level of *p* < 0.05.

To produce the full temporal profile of the relevant discriminating components (*y*(*t*)) we applied the spatial filter **w** of the window associated with the highest discrimination performance (i.e. subjected the data through the “spatial generators” leading to the most reliable discrimination) across the entire stimulus-locked window (−100 to 800 ms post-stimulus) and separately for each of the social and non-social contexts as well as the three difficulty conditions (easy, medium and difficult; Fig. 4c). The times course of these discriminating components were then z-scored separately for each participant and for each of the social and non-social contexts.

This procedure allowed us to investigate the gradual build-up of evidence accumulation activity leading up to the point of maximum discrimination and to extract the corresponding single-trial build-up rates used in subsequent analyses. These build-up rates (or slopes) were computed through a linear regression between onset and peak time of the accumulating activity extracted on a participant-specific basis. Specifically, we identified the time point at which the discriminating activity began to rise monotonically after an initial dip in the stimulus-locked data following any early evoked responses present in the data (onset time mean ± s.e.: 363.097 ± 97.046 for social trials and 376.161ms ± 107.155 for non-social trials).

Finally, the linearity of our model also allowed us to compute scalp topographies of the relevant discriminating components from Eq. 4 by estimating a forward model:

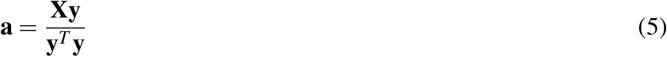

The EEG data **X** and discriminating components **y** are now depicted in matrix and vector notation, respectively, for convenience. Equation 5) represents the electrical coupling of the discriminating component **y** that explains most of the activity in **X**. Specifically, strong coupling is linked to low attenuation of the component and can be visualized as the intensity of vector **a**. We estimated forward models for the resulting discriminating activity separately for social and non-social trials (Fig. 4b).

### Single-trial regression analyses

To examine the association between the probability of reward (i.e. indirect trustworthiness levels and pure probabilities in the social and non-social contexts respectively) and the probability of playing (1: ’Play’, 0: ’Keep’) on individual trials (Fig. 3a) we performed the following single-trial logistic regression analysis (separately for each participant and for each of the social and non-social trials):

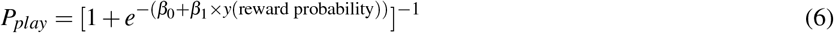

To examine the association between task difficulty (i.e. 1: easy [reward probabilities 0–0.2 and 0.8–1], 2: medium [reward probabilities 0.2–0.4 and 0.6–0.8], 3: difficult [reward probabilities 0.4–0.6]) and response times (RT) on individual trials (Fig. 3b) we performed the following single-trial regression analysis (separately for each participant and for each of the social and non-social trials):

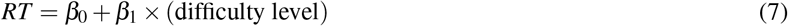

To examine the association between the rate of evidence accumulation derived from the neural data and behavioral performance on the task, we run a single-trial logistic regression analysis. Specifically, we used the trial-wise estimates of the slope of the EEG-derived evidence accumulation signal (i.e. *y*(*t*)) to predict the probability of playing (1: ’Play’, 0: ’Keep’) on individual trials (Fig. 5c, d). We performed this analysis separately for each participant and for each of the social and non-social trials:

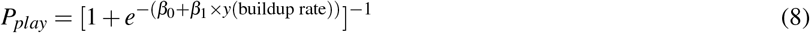

In all three cases we tested whether the regression coefficients across participants (*β*_1_ values in Eqs. 6 7 8) came from a distribution with a mean different from zero (using separate two-tailed *t* test).

### MRI data acquisition

A Siemens 3-Tesla TIM Trio MRI scanner (Siemens, Erlangen, Germany) with an 12-channel head coil was employed for the (f)MRI acquisition. A T2*-weighted gradient echo was used to acquire functional volumes with an echo-planar imaging sequence (32 interleaved slices, gap: 0.3 mm, voxel size: 3 × 3 × 3 mm, matrix size: 70 × 70, FOV: 210 mm, TE: 30 ms, TR: 2000 ms, flip angle: 80°). We recorded 5 experimental runs of 205 whole-brain volumes each, corresponding to the two blocks of trials in the main experimental task. Afterwards, we acquired phase and magnitude field maps (3 × 3 × 3 mm voxels, 32 axial slices, TR=488 ms, short TE=4.92 ms, long TE=7.38 ms) for distortion correction of the acquired EPI images. Finally, a high-resolution anatomical volume was taken using a T1-weighted sequence (192 slices, gap: 0.5 mm, voxel size: 1 × 1 × 1 mm, matrix size: 256 × 256, FOV: 256 mm, TE: 2300 ms, TR: 2.96 ms, flip angle: 9°), which was used as an anatomical reference for the functional scans.

### fMRI data preprocessing

To guarantee a steady-state fMRI we removed the first 5 volumes per run and we used only the remaining 200 volumes for the analysis. Head-related motion correction, slice-timing correction, high-pass filtering (>100s), and spatial smoothing (with a Gaussian kernel of 5mm full-width at half maximum) were performed using the FMRIB’s Software Library (Functional MRI of the Brain, Oxford, UK). The motion correction preprocessing step generated motion parameters which were subsequently included as regressors of no interest in the general linear model (GLM) analysis (see fMRI analysis below). Brain extraction of the structural and functional images was performed using the Brain Extraction tool (BET). The echo-planar imaging data for each participant was transformed into the subject-specific high-resolution space using a BBR (boundary-based registration) algorithm. The images were then registered to standard space (Montreal Neurological Institute, MNI) using FMRIB’s Non-linear Image Registration Tool with a resolution warp of 10mm and 12 degrees of freedom. Finally, to correct for signal loss and geometric distortions due to B0 field inhomogeneities B0 unwarping was used for 29 out of 31 participants. Field map images were not acquired for the remaining 2 participants.

### fMRI analysis

We performed whole-brain statistical analyses of the functional data using a multilevel approach within the framework of a GLM, as implemented in FSL (using the FEAT module) (S. M. Smith et al., 2004):

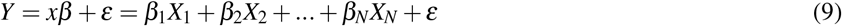

Y represents the time series (with T time samples) for a voxel and X is a T × *N* design matrix where the columns correspond to the different regressors included in the design (see below) convolved with a canonical hemodynamic response function (double-*γ* function). *β* is a *N* × 1 column vector of regression coefficients and *ε* a T × 1 column vector of residual error terms. We performed a first-level analysis to analyse each participant’s individual runs, which we then combined using a second-level analysis (fixed effects). We combined data across participants using a third-level, mixed-effects model (FLAME 1), treating participants as a random effect. Time-series statistical analysis was carried out using FMRIB’s improved linear model with local autocorrelation (Woolrich, Behrens, Beckmann, and Smith, 2005).

Our GLM included 4 regressors of interest for each of the social and non-social contexts (i.e. a total of 8 regressors). More specifically, for each of the social and non-social trials we included 1) an EEG-informed regressor with a parametric amplitude modulation based on the trial-by-trial fluctuations in the rate of evidence accumulation (i.e. trial-wise slopes in *y*(*t*)), 2) a parametric regressor with amplitude modulation based on individual trial RTs, 3) a parametric regressor with amplitude modulation based on the individual trial task difficulty (−1: difficult, 0: medium, 1: easy) and 4) an unmodulated regressor (i.e. all amplitudes set to 1) to account for any additional unaccounted variance in the data (Fig. 6a). We modelled all regressor events as boxcar functions (i.e. duration 100 ms). For the first two regressors, event onset times were aligned to the time of response, while for the last two, to the onset of stimulus presentation. Using the unmodulated regressors we also computed standard contrast and conjunction maps between social and npon-social trials. Finally, we added the motion correction parameters obtained from fMRI preprocessing (three rotations and three translations) as additional covariates of no interest.

### Resampling procedure for fMRI thresholding

In order to establish a reliable significance threshold for the fMRI data, while properly correcting for multiple comparisons, we used a resampling procedure, which examines a priori statistics of the trial-wise variability in the parametrically adjusted regressors (i.e. regressors 1–3 above) in a way that trades off cluster size and maximum voxel Z-score (Fouragnan, Retzler, Mullinger, and Philiastides, 2015; Gherman and Philiastides, 2018). Specifically, for each resampled iteration, we maintained the onset and duration of the regressors identical, whilst shuffling the amplitude values across trials, runs and participants. Thus, the resulting regressors for each participant were different as they were constructed from a random sequence of regressor amplitude events. This procedure was repeated 100 times and for each iteration we performed the full 3-level analysis (run, participant, and group). Finally, we estimated a joint threshold for the cluster size and Z-score based on the cluster outputs per shuffled regressor. This was achieved by constructing a null distribution for this joint threshold based on the size of all clusters larger than 10 voxels and with Z-scores larger than |2.57| (i.e. considering both positive and negative correlations) across all shuffled regressors. We found that the largest 5% of cluster sizes exceeded 88 voxels. We therefore used these results to derive a corrected threshold for our statistical maps, which we then applied to the clusters observed in the original data (that is, Z=±2.57, minimum cluster size of 88 voxels, corrected at p=0.05).

### Psychophysiological interaction analysis

We conducted a psychophysiological interaction (PPI) analysis to probe the functional connectivity between the pMFC found to correlate with the trial-by-trial variability in our EEG-informed regressor, and the rest of the brain. To carry out the PPI analysis, we first extracted time-series data from group-level activation clusters in the pMFC (seed), separately for each of the social and non-social contexts. Specifically, we identified the relevant pMFC clusters that were situated within the SMA and were most consistent with previous reports of EA-related activity in this region (Pisauro, Fouragnan, Retzler, and Philiastides, 2017) and then back-projected these clusters from the group (standard) space into the individual participant’s EPI (functional) space (by applying the inverse transformations estimated during the main registration procedure). The average time-series data from the back-projected voxels, which displayed activations in the direction of the predicted EA profile were then used as the physiological regressor in our PPI analysis.

The main aim of this analysis was to investigate potential task-dependant associations between the site of evidence accumulation and regions involved in domain-general value computations. If such an association exists, the coupling between these regions should be stronger while the process of evidence accumulation unfolds and it should also scale with the difficulty of the decision. To this end, our psychological regressor was constructed as a parametric boxcar regressor, the amplitude of which reflected the difficulty (1 = difficult, 2 = medium, 3 = easy) and the duration of which reflected the RT of each trial. We expected the relevant coupling to be negative, as easier trials decrease integration times and correspondingly the overall integrated activity (that is, area under the accumulation curve; Fig. 6b). The resulting fMRI statistical maps were corrected based on the threshold derived from the resampling procedure described above.

## Acknowledgements

This work was supported in part by the European Research Council (865003; MGP) and the Economic and Social Research Council (ES/L012995/1; MGP). EF was supported by a UKRI Future Leaders Fellowship (MR/T023007/1).

## Author contributions

DHA, EF, EDL and MGP conceived and designed the experiments. OGBG and PGS designed the stimuli. DHA performed the experiments and collected the data. DHA and MGP analysed the data and wrote the paper. All authors discussed the results and implications and commented on the manuscript.

## Data and code availability

The supplementary tables, data and code required to reproduce the main findings are available on: https://osf.io/hrgp5/?view_only=c84a4a66aebd4284951c8efd3a4d65fd

## References

Ahumada Jr, A., & Lovell, J. (1971). Stimulus features in signal detection. The Journal of the Acoustical Society of America, 49(6B), 1751–1756.

Basten, U., Biele, G., Heekeren, H. R., & Fiebach, C. J. (2010). How the brain integrates costs and benefits during decision making. Proceedings of the National Academy of Sciences, 107(50), 21767–21772.

Berg, D. (n.d.). Mccabe (1995) berg, j., dickhaut, j., & mccabe, k.(1995). Trust, reciprocity, and social history. Games and Economic Behavior, 10(1), 122–142.

Bogacz, R., Brown, E., Moehlis, J., Holmes, P., & Cohen, J. D. (2006). The physics of optimal decision making: A formal analysis of models of performance in two-alternative forced-choice tasks. Psychological review, 113(4), 700.

Burke, C. J., & Tobler, P. N. (2011). Coding of reward probability and risk by single neurons in animals. Frontiers in neuroscience, 5, 121.

Cerniglia, L., Bartolomeo, L., Capobianco, M., Lo Russo, S. L. M., Festucci, F., Tambelli, R., Adriani, W., & Cimino, S. (2019). Intersections and divergences between empathizing and mentalizing: Development, recent advancements by neuroimaging and the future of animal modeling. Frontiers in behavioral neuroscience, 13, 212.

Chen, F., & Krajbich, I. (2018). Biased sequential sampling underlies the effects of time pressure and delay in social decision making. Nature communications, 9(1), 1–10.

Chib, V. S., Rangel, A., Shimojo, S., & O’Doherty, J. P. (2009). Evidence for a common representation of decision values for dissimilar goods in human ventromedial prefrontal cortex. Journal of Neuroscience, 29(39), 12315–12320.

Clithero, J. A., & Rangel, A. (2014). Informatic parcellation of the network involved in the computation of subjective value. Social cognitive and affective neuroscience, 9(9), 1289–1302.

Cona, G., Marino, G., & Semenza, C. (2017). Tms of supplementary motor area (sma) facilitates mental rotation performance: Evidence for sequence processing in sma. Neuroimage, 146, 770–777.

Daw, N. D., O’doherty, J. P., Dayan, P., Seymour, B., & Dolan, R. J. (2006). Cortical substrates for exploratory decisions in humans. Nature, 441(7095), 876–879.

Domenech, P., Redouté, J., Koechlin, E., & Dreher, J.-C. (2018). The neuro-computational architecture of value-based selection in the human brain. Cerebral Cortex, 28(2), 585–601.

Donner, T. H., Siegel, M., Fries, P., & Engel, A. K. (2009). Buildup of choice-predictive activity in human motor cortex during perceptual decision making. Current Biology, 19(18), 1581–1585.

Drugowitsch, J., Moreno-Bote, R., Churchland, A. K., Shadlen, M. N., & Pouget, A. (2012). The cost of accumulating evidence in perceptual decision making. Journal of Neuroscience, 32(11), 3612–3628.

Dunbar, R. I., & Shultz, S. (2007). Evolution in the social brain. science, 317(5843), 1344–1347.

Ferrari, C., Lega, C., Vernice, M., Tamietto, M., Mende-Siedlecki, P., Vecchi, T., Todorov, A., & Cattaneo, Z. (2016). The dorsomedial prefrontal cortex plays a causal role in integrating social impressions from faces and verbal descriptions. Cerebral cortex, 26(1), 156–165.

Filimon, F., Philiastides, M. G., Nelson, J. D., Kloosterman, N. A., & Heekeren, H. R. (2013). How embodied is perceptual decision making? evidence for separate processing of perceptual and motor decisions. Journal of Neuroscience, 33(5), 2121–2136.

Forstmann, B. U., Dutilh, G., Brown, S., Neumann, J., Von Cramon, D. Y., Ridderinkhof, K. R., & Wagenmakers, E.-J. (2008). Striatum and pre-sma facilitate decision-making under time pressure. Proceedings of the National Academy of Sciences, 105(45), 17538–17542.

Fouragnan, E., Chierchia, G., Greiner, S., Neveu, R., Avesani, P., & Coricelli, G. (2013). Reputational priors magnify striatal responses to violations of trust. Journal of Neuroscience, 33(8), 3602–3611.

Fouragnan, E., Retzler, C., Mullinger, K., & Philiastides, M. G. (2015). Two spatiotemporally distinct value systems shape reward-based learning in the human brain. Nature communications, 6(1), 1–11.

Garvert, M. M., Friston, K. J., Dolan, R. J., & Garrido, M. I. (2014). Subcortical amygdala pathways enable rapid face processing. Neuroimage, 102, 309–316.

Gherman, S., & Philiastides, M. G. (2015). Neural representations of confidence emerge from the process of decision formation during perceptual choices. Neuroimage, 106, 134–143.

Gherman, S., & Philiastides, M. G. (2018). Human vmpfc encodes early signatures of confidence in perceptual decisions. Elife, 7, e38293.

Gill, D., Garrod, O. G., Jack, R. E., & Schyns, P. G. (2014). Facial movements strategically camouflage involuntary social signals of face morphology. Psychological science, 25(5), 1079–1086.

Gluth, S., Hotaling, J. M., & Rieskamp, J. (2017). The attraction effect modulates reward prediction errors and intertemporal choices. Journal of Neuroscience, 37(2), 371–382.

Gluth, S., Rieskamp, J., & Büchel, C. (2012). Deciding when to decide: Time-variant sequential sampling models explain the emergence of value-based decisions in the human brain. Journal of Neuroscience, 32(31), 10686–10698.

Griessinger, T., & Coricelli, G. (2015). The neuroeconomics of strategic interaction. Current Opinion in Behavioral Sciences, 3, 73–79.

Grinband, J., Wager, T. D., Lindquist, M., Ferrera, V. P., & Hirsch, J. (2008). Detection of time-varying signals in event-related fmri designs. Neuroimage, 43(3), 509–520.

Hare, T. A., Schultz, W., Camerer, C. F., O’Doherty, J. P., & Rangel, A. (2011). Transformation of stimulus value signals into motor commands during simple choice. Proceedings of the National Academy of Sciences, 108(44), 18120–18125.

Herding, J., Ludwig, S., von Lautz, A., Spitzer, B., & Blankenburg, F. (2019). Centro-parietal eeg potentials index subjective evidence and confidence during perceptual decision making. NeuroImage, 201, 116011.

Hoel, P., Port, S., & Stone, C. (1971). Introduction to probability theory. series in statistics.

Hu, C., Domenech, P., & Pessiglione, M. (2020). Order matters: How covert value updating during sequential option sampling shapes economic preference. PLoS computational biology, 16(8), e1007920.

Hunt, L. T., Kolling, N., Soltani, A., Woolrich, M. W., Rushworth, M. F., & Behrens, T. E. (2012). Mechanisms underlying cortical activity during value-guided choice. Nature neuroscience, 15(3), 470–476.

Hutcherson, C. A., Montaser-Kouhsari, L., Woodward, J., & Rangel, A. (2015). Emotional and utilitarian appraisals of moral dilemmas are encoded in separate areas and integrated in ventromedial prefrontal cortex. Journal of Neuroscience, 35(36), 12593–12605.

Ivanoff, J., Branning, P., & Marois, R. (2008). Fmri evidence for a dual process account of the speed-accuracy tradeoff in decision-making. PLoS one, 3(7), e2635.

Kelly, S. P., & O’Connell, R. G. (2013). Internal and external influences on the rate of sensory evidence accumulation in the human brain. Journal of Neuroscience, 33(50), 19434–19441.

Kim, J.-H., Lee, J.-M., Jo, H. J., Kim, S. H., Lee, J. H., Kim, S. T., Seo, S. W., Cox, R. W., Na, D. L., Kim, S. I., et al. (2010). Defining functional sma and pre-sma subregions in human mfc using resting state fmri: Functional connectivity-based parcellation method. Neuroimage, 49(3), 2375–2386.

Kolling, N., Braunsdorf, M., Vijayakumar, S., Bekkering, H., Toni, I., & Mars, R. B. (2021). Constructing others’ beliefs from one’s own using medial frontal cortex. Journal of Neuroscience.

Krajbich, I., Hare, T., Bartling, B., Morishima, Y., & Fehr, E. (2015). A common mechanism underlying food choice and social decisions. PLoS Comput Biol, 11(10), e1004371.

Lieberman, M. D., Straccia, M. A., Meyer, M. L., Du, M., & Tan, K. M. (2019). Social, self,(situational), and affective processes in medial prefrontal cortex (mpfc): Causal, multivariate, and reverse inference evidence. Neuroscience & Biobehavioral Reviews, 99, 311–328.

Lim, S.-L., O’Doherty, J. P., & Rangel, A. (2013). Stimulus value signals in ventromedial pfc reflect the integration of attribute value signals computed in fusiform gyrus and posterior superior temporal gyrus. Journal of Neuroscience, 33(20), 8729–8741.

Liu, T., & Pleskac, T. J. (2011). Neural correlates of evidence accumulation in a perceptual decision task. Journal of neurophysiology, 106(5), 2383–2398.

Lockwood, P. L., Apps, M. A., & Chang, S. W. (2020). Is there a ‘social’brain? implementations and algorithms. Trends in Cognitive Sciences.

Lockwood, P. L., Hamonet, M., Zhang, S. H., Ratnavel, A., Salmony, F. U., Husain, M., & Apps, M. A. (2017). Prosocial apathy for helping others when effort is required. Nature human behaviour, 1(7), 0131.

Monosov, I. E. (2017). Anterior cingulate is a source of valence-specific information about value and uncertainty. Nature communications, 8(1), 1–12.

Morgenstern, O., & Von Neumann, J. (1953). Theory of games and economic behavior. Princeton university press.

Mulder, M., Van Maanen, L., & Forstmann, B. (2014). Perceptual decision neurosciences–a model-based review. Neuroscience, 277, 872–884.

Mumford, J. A., Poline, J.-B., & Poldrack, R. A. (2015). Orthogonalization of regressors in fmri models. PloS one, 10(4), e0126255.

Nachev, P., Kennard, C., & Husain, M. (2008). Functional role of the supplementary and pre-supplementary motor areas. Nature Reviews Neuroscience, 9(11), 856–869.

Neubert, F.-X., Mars, R. B., Sallet, J., & Rushworth, M. F. (2015). Connectivity reveals relationship of brain areas for reward-guided learning and decision making in human and monkey frontal cortex. Proceedings of the national academy of sciences, 112(20), E2695–E2704.

Parra, L. C., Spence, C. D., Gerson, A. D., & Sajda, P. (2005). Recipes for the linear analysis of eeg. Neuroimage, 28(2), 326–341.

Pearson, J. M., Watson, K. K., & Platt, M. L. (2014). Decision making: The neuroethological turn. Neuron, 82(5), 950–965.

Peixoto, D., Verhein, J. R., Kiani, R., Kao, J. C., Nuyujukian, P., Chandrasekaran, C., Brown, J., Fong, S., Ryu, S. I., Shenoy, K. V., et al. (2021). Decoding and perturbing decision states in real time. Nature, 591(7851), 604–609.

Philiastides, M. G., Biele, G., & Heekeren, H. R. (2010). A mechanistic account of value computation in the human brain. Proceedings of the National Academy of Sciences, 107(20), 9430–9435.

Philiastides, M. G., Heekeren, H. R., & Sajda, P. (2014). Human scalp potentials reflect a mixture of decision-related signals during perceptual choices. Journal of Neuroscience, 34(50), 16877–16889.

Philiastides, M. G., & Sajda, P. (2006). Temporal characterization of the neural correlates of perceptual decision making in the human brain. Cerebral cortex, 16(4), 509–518.

Philiastides, M. G., & Sajda, P. (2007). Eeg-informed fmri reveals spatiotemporal characteristics of perceptual decision making. Journal of Neuroscience, 27(48), 13082–13091.

Pisauro, M. A., Fouragnan, E., Retzler, C., & Philiastides, M. G. (2017). Neural correlates of evidence accumulation during value-based decisions revealed via simultaneous eeg-fmri. Nature communications, 8(1), 1–9.

Piva, M., Velnoskey, K., Jia, R., Nair, A., Levy, I., & Chang, S. W. (2019). The dorsomedial prefrontal cortex computes task-invariant relative subjective value for self and other. Elife, 8, e44939.

Polaniía, R., Krajbich, I., Grueschow, M., & Ruff, C. C. (2014). Neural oscillations and synchronization differentially support evidence accumulation in perceptual and value-based decision making. Neuron, 82(3), 709–720.

Rangel, A., & Hare, T. (2010). Neural computations associated with goal-directed choice. Current opinion in neurobiology, 20(2), 262–270.

Ratcliff, R., Philiastides, M. G., & Sajda, P. (2009). Quality of evidence for perceptual decision making is indexed by trial-to-trial variability of the eeg. Proceedings of the National Academy of Sciences, 106(16), 6539–6544.

Rilling, J. K., King-Casas, B., & Sanfey, A. G. (2008). The neurobiology of social decision-making. Current opinion in neurobiology, 18(2), 159–165.

Ruff, C. C., & Fehr, E. (2014). The neurobiology of rewards and values in social decision making. Nature Reviews Neuroscience, 15(8), 549–562.

Sajda, P., Philiastides, M. G., Heekeren, H., & Ratcliff, R. (2011). Linking neuronal variability to perceptual decision making via neuroimaging. In M. Ding & D. Glanzman (Eds.), The dynamic brain: An exploration of neuronal variability and its functional significance (pp. 214–232). Oxford University Press.

Sajda, P., Philiastides, M. G., & Parra, L. C. (2009). Single-trial analysis of neuroimaging data: Inferring neural networks underlying perceptual decision-making in the human brain. IEEE reviews in biomedical engineering, 2, 97–109.

Scott, L. S., & Monesson, A. (2009). The origin of biases in face perception. Psychological Science, 20(6), 676–680.

Sepulveda, P., Usher, M., Davies, N., Benson, A. A., Ortoleva, P., & De Martino, B. (2020). Visual attention modulates the integration of goal-relevant evidence and not value. Elife, 9, e60705.

Skelly, L. R., & Decety, J. (2012). Passive and motivated perception of emotional faces: Qualitative and quantitative changes in the face processing network. PloS one, 7(6), e40371.

Smith, B. W., Mitchell, D. G., Hardin, M. G., Jazbec, S., Fridberg, D., Blair, R. J. R., & Ernst, M. (2009). Neural substrates of reward magnitude, probability, and risk during a wheel of fortune decision-making task. Neuroimage, 44(2), 600–609.

Smith, D. V., Hayden, B. Y., Truong, T.-K., Song, A. W., Platt, M. L., & Huettel, S. A. (2010). Distinct value signals in anterior and posterior ventromedial prefrontal cortex. Journal of Neuroscience, 30(7), 2490–2495.

Smith, S. M., Jenkinson, M., Woolrich, M. W., Beckmann, C. F., Behrens, T. E., Johansen-Berg, H., Bannister, P. R., De Luca, M., Drobnjak, I., Flitney, D. E., et al. (2004). Advances in functional and structural mr image analysis and implementation as fsl. Neuroimage, 23, S208–S219.

Steinemann, N. A., O’Connell, R. G., & Kelly, S. P. (2018). Decisions are expedited through multiple neural adjustments spanning the sensorimotor hierarchy. Nature communications, 9(1), 1–13.

Sugrue, L. P., Corrado, G. S., & Newsome, W. T. (2004). Matching behavior and the representation of value in the parietal cortex. science, 304(5678), 1782–1787.

Suzuki, S., & O’Doherty, J. P. (2020). Breaking human social decision making into multiple components and then putting them together again. Cortex.

Turner, B. M., Van Maanen, L., & Forstmann, B. U. (2015). Informing cognitive abstractions through neuroimaging: The neural drift diffusion model. Psychological review, 122(2), 312.

Uleman, J. S., & Kressel, L. M. (2013). A brief history of theory and research on impression formation.

Van Overwalle, F. (2009). Social cognition and the brain: A meta-analysis. Human brain mapping, 30(3), 829–858.

van Baar, J. M., Chang, L. J., & Sanfey, A. G. (2019). The computational and neural substrates of moral strategies in social decision-making. Nature communications, 10(1), 1–14.

van Vugt, M. K., Beulen, M. A., & Taatgen, N. A. (2019). Relation between centro-parietal positivity and diffusion model parameters in both perceptual and memory-based decision making. Brain research, 1715, 1–12.

Verdonck, S., Loossens, T., & Philiastides, M. G. (2021). The leaky integrating threshold and its impact on evidence accumulation models of choice rt. Psychological Review, 128(2), 203–221.

Voss, A., Rothermund, K., & Voss, J. (2004). Interpreting the parameters of the diffusion model: An empirical validation. Memory & cognition, 32(7), 1206–1220.

Vuilleumier, P., Richardson, M. P., Armony, J. L., Driver, J., & Dolan, R. J. (2004). Distant influences of amygdala lesion on visual cortical activation during emotional face processing. Nature neuroscience, 7(11), 1271–1278.

Woolrich, M. W., Behrens, T. E. J., Beckmann, C. F., & Smith, S. M. (2005). Mixture models with adaptive spatial regularization for segmentation with an application to fmri data. IEEE transactions on medical imaging, 24(1), 1–11.

Wu, S.-W., Delgado, M. R., & Maloney, L. T. (2015). Gambling on visual performance: Neural correlates of metacognitive choice between visual lotteries. Frontiers in neuroscience, 9, 314.

Wu, Z., Litwin-Kumar, A., Shamash, P., Taylor, A., Axel, R., & Shadlen, M. N. (2020). Context-dependent decision making in a premotor circuit. Neuron, 106(2), 316–328.

Yu, H., Garrod, O. G., & Schyns, P. G. (2012). Perception-driven facial expression synthesis. Computers & Graphics, 36(3), 152–162.

Zhan, J., Garrod, O. G., van Rijsbergen, N., & Schyns, P. G. (2019). Modelling face memory reveals task-generalizable representations. Nature human behaviour, 3(8), 817–826.

Zhang, J., Hughes, L. E., & Rowe, J. B. (2012). Selection and inhibition mechanisms for human voluntary action decisions. Neuroimage, 63(1), 392–402.

